# Multi-dimensional demarcation of phylogenetic groups of plant 14-3-3 isoforms using biochemical signatures

**DOI:** 10.1101/2024.07.02.601760

**Authors:** Ilya A. Sedlov, Nikolai N. Sluchanko

## Abstract

Interaction of dimeric 14-3-3 proteins with numerous phosphotargets regulates various physiological processes in plants, from flowering to transpiration and salt tolerance. Several genes express distinct 14-3-3 ‘isoforms’, particularly numerous in plants, but comparative studies of all 14-3-3 isoforms for a given organism have not been undertaken. Here we systematically investigated twelve 14-3-3 isoforms from the model plant *Arabidopsis thaliana*, uniformly capable of homodimerization at high protein concentration. We unexpectedly discovered that, at physiological protein concentrations, four isoforms representing a seemingly more ancestral, epsilon phylogenetic group (iota, mu, omicron, epsilon) demonstrate an outstanding monomerization propensity and enhanced surface hydrophobicity, which is uncharacteristic for eight non-epsilon isoforms (omega, phi, chi, psi, upsilon, nu, kappa, lambda). Further analysis revealed that dramatically lowered thermodynamic stabilities entail aggregation of the epsilon-group isoforms at near-physiological temperatures and provoke their proteolytic degradation. Structure-inspired single mutations in 14-3-3 iota could rescue non-epsilon behavior, thereby pinpointing key positions responsible for the phylogenetic demarcation. Combining two major demarcating positions (namely, 27th and 51st in omega) and multi-dimensional differences in biochemical properties identified here, we developed a predictor strongly supporting categorization of abundant 14-3-3 isoforms widely across plant groups, from Eudicots to Monocots, Gymnosperms and Lycophytes. In particular, our approach fully recapitulates the phylogenetic epsilon/non-epsilon demarcation in Eudicots and supports the presence of isoforms of both types in more primitive plant groups such as *Selaginella*, thereby refining solely sequence-based analysis in evolutionarily distant species and providing novel insights into the evolutionary history of the epsilon phylogenetic group.

**Significance:** Despite over 30 years of research, systematic comparative studies on the regulatory plant 14-3-3 proteins have not been undertaken, making phylogenetic classification of numerous plant 14-3-3 isoforms in different species unreliable. Working on twelve purified *Arabidopsis* 14-3-3 isoforms, we have discovered a set of biochemical signatures that can be used to robustly and widely categorize epsilon and non-epsilon plant 14-3-3 isoforms, also identifying at least two amino acid positions responsible for such multi-dimensional demarcation.

## Introduction

The study of 14-3-3 proteins has more than 60 years of history (Moore and Perez, 1967). A total of seven 14-3-3 isoforms were found in mammals, they were called according to some of the first letters of the Greek alphabet (β, γ, ε, ζ, η, τ, σ) (Isobe *et al*., 1991). 14-3-3s bind and regulate phosphorylated protein partners, influencing activity and stability of the latter (Muslin *et al*., 1996). Acting as adaptors and regulators of client protein activity, subcellular localization and interaction with other cell factors, 14-3-3s contribute to multiple physiological processes (Mackintosh, 2004; Aitken, 2006; Pennington *et al*., 2018; van Heusden, 2005; Denison *et al*., 2011). Nitrate reductase (Moorhead *et al*., 1996) and plasma membrane H^+^ ATP-ase (Ottmann *et al*., 2007) have become classical examples of plant phosphoproteins regulated by 14-3-3, whereas the 14-3-3 role in regulating photoperiodism (Mayfield *et al*., 2007), flowering (Taoka *et al*., 2011), root growth and chloroplast development (Mayfield *et al*., 2012) as well as in responses to salt and other stress (XU and SHI, 2006) is well-established. These proteins are small and acidic, and normally possess an alpha-helical W-shaped dimeric structure with a single phosphopeptide-binding site located in each subunit (Gardino *et al*., 2006; Yang *et al*., 2006; Liu *et al*., 1995; Yaffe *et al*., 1997) (Fig. 1A). Dimeric status is believed to be important for their phosphopeptide-binding functions; however, some 14-3-3s monomerize as a consequence of specific phosphorylation at residues in the dimeric interface and may have specific functions such as anti-aggregating activity toward misfolded client proteins (Woodcock *et al*., 2003; Denison *et al*., 2014; Sluchanko *et al*., 2008; Trošanová *et al*., 2022; Gu *et al*., 2006; Zhu *et al*., 2023).

**Fig. 1.**
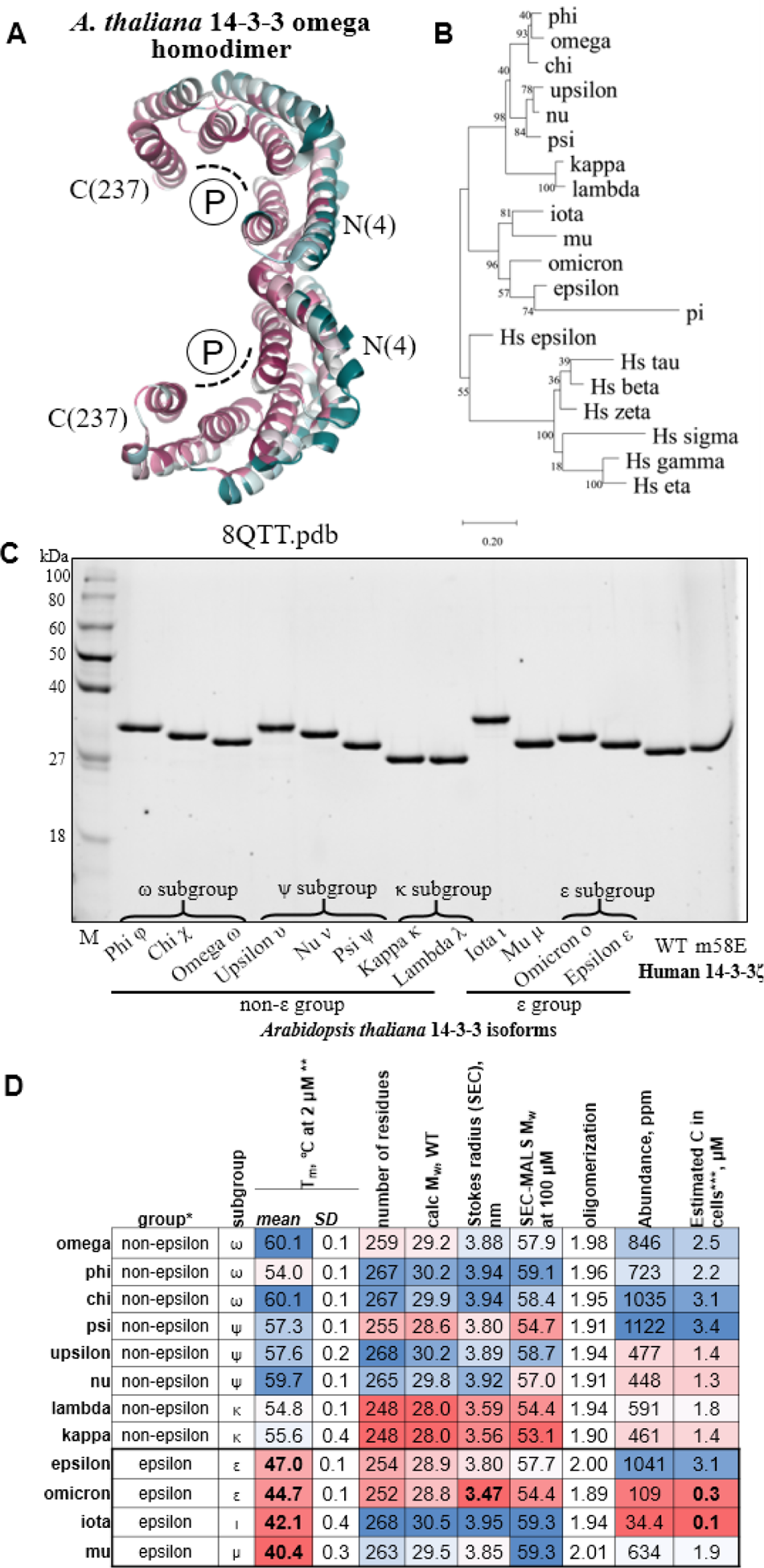
*Arabidopsis thaliana* 14-3-3 isoforms. **A**. Crystal structure of *A. thaliana* 14-3-3 omega colored according to standard Consurf (Ashkenazy *et al*., 2016) conservation levels from high (purple) to low (cyan). For Consurf analysis, 150 nonredundant homologs with 50-95% identity to *A. thaliana* 14-3-3 omega were automatically selected, aligned and scored. Dashed lines show the location of two conserved phosphopeptide-binding regions (P). **B**. Phylogenetic tree showing the main groups and subgroups of plant 14-3-3 isoforms built using the seven human 14-3-3 isoforms as an outgroup, according to the maximum likelihood method in MEGA11 using bootstrap randomization of 1000 (Tamura *et al*., 2021). Numbers indicate percentage of bootstrap operations that led to the topology shown. The scale shows evolutionary distance. **C**. SDS-PAGE analysis of 12 recombinantly produced isoforms of *A. thaliana* and two versions of human 14-3-3 zeta (ζ): the wild-type and a monomeric mutant ζm58E (Sluchanko and Uversky, 2015). 0.6 μg of each protein was loaded on the gel. **D**. Properties of twelve 14-3-3 isoforms from *A. thaliana*. The colored columns are represented by a conditional formatting tool in Microsoft Excel using a color gradient from low values (0%, red), to intermediate values (50%, white), to high values (100%, blue), within each column independently. The epsilon group isoforms are outlined by a bold line for convenience of comparisons and finding correlations. *phylogenetics according to (Mikhaylova *et al*., 2021). **for epsilon the value for 5 μM is shown. Mean ± standard deviation (*n*=3) are shown for each isoform. ***calculated given the intracellular concentration of all proteins equal to 3 mM (Milo, 2013) and abundances of *Arabidopsis* 14-3-3 isoforms taken from PaxDB (Wang *et al*., 2015).

Plant 14-3-3s were first characterized in the early 1990s due to homology with mammalian proteins (de Vetten *et al*., 1992; Guihua Lu *et al*., 1994; G Lu *et al*., 1994; Daugherty *et al*., 1996) and designated as GRF (<u>G</u>eneral <u>R</u>egulatory <u>F</u>actor) or GF14 (<u>G</u>-box <u>F</u>actor 14-3-3). In a model plant *Arabidopsis thaliana* (mouse-ear cress), 13 known genes of 14-3-3, like mammalian ones, were denoted according to the Greek alphabet, but starting from its end: omega (ω), psi (ψ), chi (χ), phi (φ), upsilon (υ), pi (π), omicron (ο), nu (ν), mu (μ), lambda (λ), kappa (κ), iota (ι), epsilon (ε) (Fig. 1B). This “*Arabidopsis*-based” classification with the Greek letter designations is frequently applied to orthologous proteins in other plant species (Mikhaylova *et al*., 2021; Hernández-Domínguez *et al*., 2019).

The total number of 14-3-3 isoforms vary between plant species, e.g., from 5 in strawberry (*Fragaria vesca*) (Mikhaylova *et al*., 2021), 8 in rice (*Oryza sativa*) (Yao *et al*., 2007), 13 in tomato (*Solanum lycopersicum*) (Jia *et al*., 2022), 18 in soybean (*Glycine max*) (Li and Dhaubhadel, 2011), 26 in tea (*Camellia sinensis*) (Zhang *et al*., 2022), to 36 in apple (*Malus domestica*) (Ren *et al*., 2023). Similar to other plant genes, the 14-3-3 diversity in plants is a result of segmental and whole genome duplications, accompanying plant genomes through evolution (Landis *et al*., 2018; Carretero-Paulet and Fares, 2012; Cao and Yan, 2016; Zuo *et al*., 2021; Ren *et al*., 2023). Multiple evolutionary analyses indicated the fundamental divergence event which led to the separation of two large phylogenetic clades called epsilon- and non-epsilon groups, however, their biochemical and functional distinction remain incompletely understood (Mikhaylova *et al*., 2021; Ren *et al*., 2023; Cao and Yan, 2016; Zuo *et al*., 2021). Epsilon-group isoforms and their conserved functions in the plant organism are sometimes considered ancestral (Keicher *et al*., 2017), which would be partly supported by their sole presence in primitive plant groups, however this remains questionable especially given large evolutionary distances and uncertainty of solely phylogenetic analysis (Cao and Yan, 2016; Zhang *et al*., 2022).

Inside the epsilon and non-epsilon groups, the diversification process went further to produce subgroups (Mikhaylova *et al*., 2021; Sehnke *et al*., 2002) (Fig. 1C and D). In *A. thaliana*, there are 5 proteins in the epsilon-group (1 iota (ι), 1 mu (μ) and 3 epsilon (ο, π, ε) isoforms) and 8 proteins in the non-epsilon group (3 omega (χ, φ, ω), 3 psi (υ, ν, ψ) and 2 kappa (κ, λ) isoforms), with pi being the most divergent isoform (Supplementary Fig. 1) expressed in extremely low amounts (Keicher *et al*., 2017). Notably, not all subgroups are represented in every plant taxa. For example, the epsilon subgroup is absent in *Fabaceae*. In *Poales*, a single epsilon-group isoform is present and three subgroups (iota, mu, kappa) are missing (Mikhaylova *et al*., 2021).

Although functions of different 14-3-3 isoforms apparently overlap, a number of studies have demonstrated functions specific to either phylogenetic group. For instance, out of eight tested *A. thaliana* 14-3-3 isoforms, omega, kappa and lambda (non-epsilon group) were the strongest inhibitors of the nitrate reductase, while 14-3-3 epsilon had almost no inhibitory effect (Lambeck *et al*., 2010). Another study demonstrated that in the case of *A. thaliana* 14-3-3 mu (epsilon-group), and not 14-3-3 upsilon (non-epsilon group), a loss-of-function mutation is associated with a decreased root length under normal light (Mayfield *et al*., 2012). Swatek et al. revealed that, while both 14-3-3 chi (non-epsilon group) and 14-3-3 epsilon interact with client proteins involved in various metabolic pathways, including glycolysis and *de novo* fatty acid synthesis, their client binding preferences are different, which illustrates 14-3-3 isoform specificity in *Arabidopsis* seed development (Swatek *et al*., 2011). Expression of 14-3-3 isoforms varies in different tissues and organs (Keicher *et al*., 2017; Rosenquist *et al*., 2001; Wilson *et al*., 2016) (Fig. 1D), which likely supports specialized functions.

Different sets of 14-3-3 isoforms in plant species require reliable biochemical/functional categorization, yet the phylogenetic analysis based on entire amino acid sequences of 14-3-3 isoforms becomes uncertain for distant species. Furthermore, the lack of a comprehensive comparison of all 14-3-3 isoforms even for the model organism, *A. thaliana*, substantially limits mechanistic interpretation of multiple observations made on a cellular, tissue and organismal level.

To fill these gaps, here we heterologously expressed all thirteen *Arabidopsis* 14-3-3 isoforms and examined twelve isoforms except pi, which could not be obtained in a soluble form, by a combination of biochemical and biophysical techniques. This helped us derive a set of signatures for robust multi-dimensional, data-based demarcation of epsilon and non-epsilon 14-3-3 groups across a wide range of plant groups. The results were validated by bioinformatics analysis, data mining in databases and by site-directed mutagenesis.

## Results

### Twelve *A. thaliana* 14-3-3 isoforms form homodimers differing by the apparent size

Out of 13 canonical isoforms encoded in *A. thaliana* we managed to obtain 12 (Fig. 1D and Supplementary Fig. 1), whereas all our attempts to obtain soluble 14-3-3 pi have been unsuccessful even in the presence of the known solubility enhancer, maltose-binding protein (MBP) (Sluchanko *et al*., 2016; Ko *et al*., 2021) (Supplementary Fig. 2). Twelve other 14-3-3 isoforms representing two major phylogenetic groups of plant 14-3-3 proteins, i.e., epsilon (epsilon, iota, omicron, mu) and non-epsilon (chi, omega, phi, upsilon, nu, psi, kappa, lambda) could be purified to homogeneity (Fig. 1C). Trypsinolysis and subsequent MALDI-MS revealed >90% sequence coverage and intactness of the terminal regions for all 12 proteins studied (Supplementary Fig. 1), whereas the electrophoretic mobility pattern on SDS-PAGE was consistent with the the expected sizes of the polypeptide chains (Fig. 1C and D). We then used affinity-purified protein samples before SEC, to detect all possible oligomeric forms.

An absolute molecular mass analysis using SEC-MALS at 100 μM revealed major symmetrical peaks for all twelve 14-3-3 isoforms from *A. thaliana* with *M*_w_ values closely matching the expected dimeric masses (‘oligomerization’ was equal to 1.89-2.01) (Fig. 1D and Supplementary Fig. 3). While the MALS-derived dimeric masses for all twelve isoforms were close to the calculated values (Fig. 1D), we noticed a significant variation of the apparent size of these dimers reflected in the Stokes radii ranging from 3.95 nm (iota) to 3.47 nm (omicron). Although iota (calculated dimeric *M*_w_ 61 kDa) was larger than omicron (calculated dimeric *M*_w_ 57.6 kDa) by only 3.4 kDa, the apparent *M*_w_ values determined from SEC peak position corresponded to ∼85 and ∼60 kDa, respectively, creating a 25 kDa difference.

### All epsilon-group 14-3-3 homodimers exhibit concentration-dependent monomerization

The differences in the apparent size of 14-3-3 proteins may reflect a significant contribution from their variable C-terminal tails. Indeed, these elements vary in length, sequences and conformation (Supplementary Fig. 1) (W., Shen *et al*., 2003; Truong *et al*., 2002). Another possibility was that the isoforms showing the latest elution on SEC were partially dissociated. To exclude this, we subjected all twelve *Arabidopsis* isoforms to gel electrophoresis under non-denaturing conditions, along with human 14-3-3 epsilon, 14-3-3 zeta and its monomerized variant (Sluchanko and Uversky, 2015) used as controls. Starting from this experiment, all 14-3-3 samples were additionally purified by preparative SEC, where the major dimeric peaks were collected and concentrated for further studies.

On native PAGE (Fig. 2A), eight non-epsilon *Arabidopsis* 14-3-3 isoforms migrated as sharp bands with the mobility similar or lower than that of dimeric human 14-3-3 zeta, and, therefore, apparently represented dimers with the electrophoretic mobilities reflecting differences in the polypeptide lengths (Fig. 1D). Totally unexpectedly, all four epsilon group *Arabidopsis* 14-3-3 isoforms (iota, mu, epsilon and, to a lesser extent, omicron), showed a downward smearing pattern indicative of monomerization. This was particularly clear from the comparison with the electrophoretic mobilities of the dimeric and monomeric 14-3-3 zeta samples used as controls (Fig. 2A). A similar smearing pattern was exhibited by human 14-3-3 epsilon, the most divergent human isoform located on the phylogenetic tree closest to *Arabidopsis* 14-3-3 proteins (Fig. 1B), and by phosphomimicking mutants of mammalian and plant 14-3-3 isoforms (Sluchanko *et al*., 2008; Denison *et al*., 2014). Since spontaneous phosphorylation of target proteins is documented in *E. coli* (Brokx *et al*., 2011; Bose *et al*., 1999), we excluded that the observed smearing electrophoretic pattern results from such modification by subjecting *Arabidopsis* 14-3-3 omega, iota, mu, omicron, epsilon and human 14-3-3 epsilon to dephosphorylation by unspecific lambda phosphatase (Supplementary Fig. 4) (Tugaeva *et al*., 2023).

**Fig. 2.**
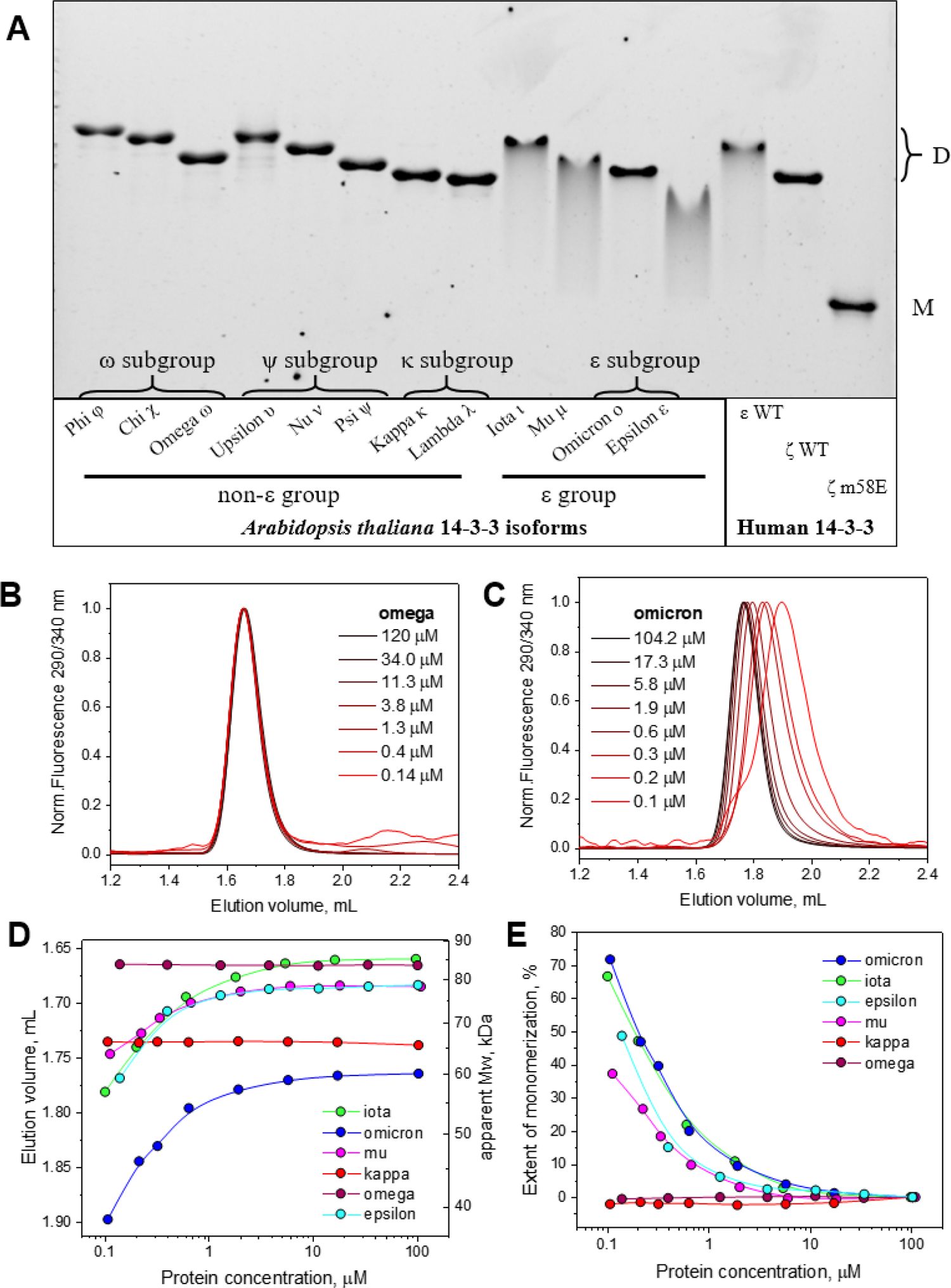
Epsilon group-specific 14-3-3 dimer dissociation. **A**. Dimer dissociation of epsilon-group *A. thaliana* 14-3-3 isoforms revealed by native PAGE. Samples of human 14-3-3 epsilon wild-type (ε WT), 14-3-3 zeta wild-type (ζ WT) and the monomerized mutant form of the latter (ζm58E) (Sluchanko and Uversky, 2015) were used for comparison. Positions of dimers (D) and monomers (M) are shown. 0.6 μg of each protein was loaded on the gel with a 15-25% acrylamide gradient, run for 2 h at room temperature and visualized by Stain-Free technology (Bio-Rad). Note the different electrophoretic mobilities of various dimeric 14-3-3 isoforms. **B, C**. Analytical SEC profiles of 14-3-3 omega (B) and omicron (C) at different protein concentrations in the loaded sample (indicated in μM). Elution profiles from a Superdex 200 Increase 5/150 column (0.45 ml/min) were followed by Trp fluorescence detection (excitation 290 nm, emission 340 nm). **D**. Concentration-dependent changes of the elution volumes determined at the peak maxima on the elution profiles of studied 14-3-3 isoforms. Right Y axis shows the apparent Mw values determined for the elution volumes of 14-3-3 species from calibrating the column with standard proteins. Note that the right and bottom axes are logarithmic. **E**. The extent of the dimer-to-monomer transition for each of the six isoforms presented on panel D.

To confirm that monomerization is not limited to the conditions of electrophoresis, we subjected two representatives of non-epsilon 14-3-3 (omega and kappa) and all four epsilon-group isoforms (iota, omicron, mu, epsilon) to analytical SEC at varying protein concentrations (Fig. 2B, C and Supplementary Fig. 5). Fluorescence-assisted detection allowed us to observe protein elution peaks even at extremely low protein concentrations (nM). In this experiment, both non-epsilon 14-3-3 isoforms showed an unchanged position on the elution profile, in a range of protein concentrations spanning three orders of magnitude (∼100 nM–100 μM). In striking contrast, all four epsilon 14-3-3 isoforms demonstrated concentration-dependent shifts toward the larger elution volumes, indicating monomerization (Fig. 2B, C and Supplementary Fig. 5). Plotting the dependence of the elution volume (or the apparent *M*_w_) on protein concentration has allowed us to assess the magnitude of the monomerization effect better (Fig. 2D). Assuming that the apparent *M*_w_ at 100 μM for each tested isoform corresponds to its fully dimeric state and that the apparent *M*_w_ of its monomer is twice as smaller, we were able to estimate the degree of dimer-to-monomer transition at each protein concentration analyzed (Fig. 2E). This plot clearly demonstrates that only epsilon-group isoforms have the monomerization propensity and that at lowest concentrations analyzed this can reach as high as ∼70 % (for iota and omicron).

That monomerization of epsilon-group 14-3-3 isoforms is the essence of the observed phenomenon was directly confirmed via forced dimerization by chemical crosslinking with bis(sulfosuccinimidyl) suberate (BS3), alongside with omega and two human 14-3-3 isoforms (zeta and epsilon) as controls (Supplementary Fig. 6).

### Monomerized epsilon-group 14-3-3 isoforms have enhanced surface hydrophobicity

Monomerization of 14-3-3 is usually accompanied with a substantial increase of surface hydrophobicity due to the exposure of hydrophobic residues from the dimerization interface (Sluchanko *et al*., 2014; Sluchanko *et al*., 2011; Sluchanko and Gusev, 2017; Woodcock *et al*., 2017). We therefore compared the hydrophobic properties of *Arabidopsis* 14-3-3 isoforms (2 μM) by their ability to interact with bis-ANS (10 μM). Bis-ANS is a solvatofluorochromic dye having a very low quantum yield of fluorescence in aqueous buffer but high green fluorescence in nonpolar environments or when bound by protein hydrophobic regions (Sluchanko *et al*., 2008; Sluchanko *et al*., 2014). Under the experimental conditions used, we observed dramatically enhanced bis-ANS fluorescence (∼6 times on average) for the four 14-3-3 isoforms of the epsilon phylogenetic group (iota, omicron, epsilon and mu) compared with the non-epsilon isoforms (Supplementary Fig. 7).

### Outstanding instability of the epsilon-group 14-3-3 isoforms

Since 14-3-3 isoforms of the epsilon and non-epsilon groups differed by stability of their homodimers, we systematically analyzed them by thermal shift assays. Fig. 3A shows that all twelve 14-3-3 isoforms from *A. thaliana* display the heat-induced denaturation profile with a single transition occurring at surprisingly different temperatures – from ∼40.4 °C in the case of mu to 60.1 °C in the case of omega and chi. The epsilon group isoforms exhibited much lower *T*_m_ values (43.6 °C on average) compared to the non-epsilon group isoforms (57.4 °C on average). Of note, the observed *T*_m_ values within the epsilon group *Arabidopsis* isoforms were significantly lower than even the *T*_m_ of the human 14-3-3 zeta monomer (52.2°C), whereas the *T*_m_ values for most of the non-epsilon group *Arabidopsis* isoforms were similar to those of dimeric human 14-3-3 zeta (Fig. 3A). With the caveat of appreciable variability of *T*_m_ within the groups, the denaturation profiles of the non-epsilon group isoforms most probably corresponded to denaturation of the dimeric 14-3-3, whereas the profiles of the epsilon-group isoforms corresponded to denaturation of the largely monomerized 14-3-3. In agreement with this notion, we observed a concentration-dependent change of the thermal stability profile for *Arabidopsis* 14-3-3 epsilon (Supplementary Fig. 8). To exclude that the lower *T*_m_ for the epsilon-group isoforms is a consequence of peculiar properties of the thermal shift method used (e.g., a hydrophobic ProteOrange dye-induced 14-3-3 monomerization), we used an alternative assay based on measuring intrinsic tryptophan fluorescence. This method showed nearly identical results indicating that iota is almost 20 °C less stable than omega or human 14-3-3 zeta (Supplementary Fig. 8), thereby verifying the data obtained using ProteOrange dye.

**Fig. 3.**
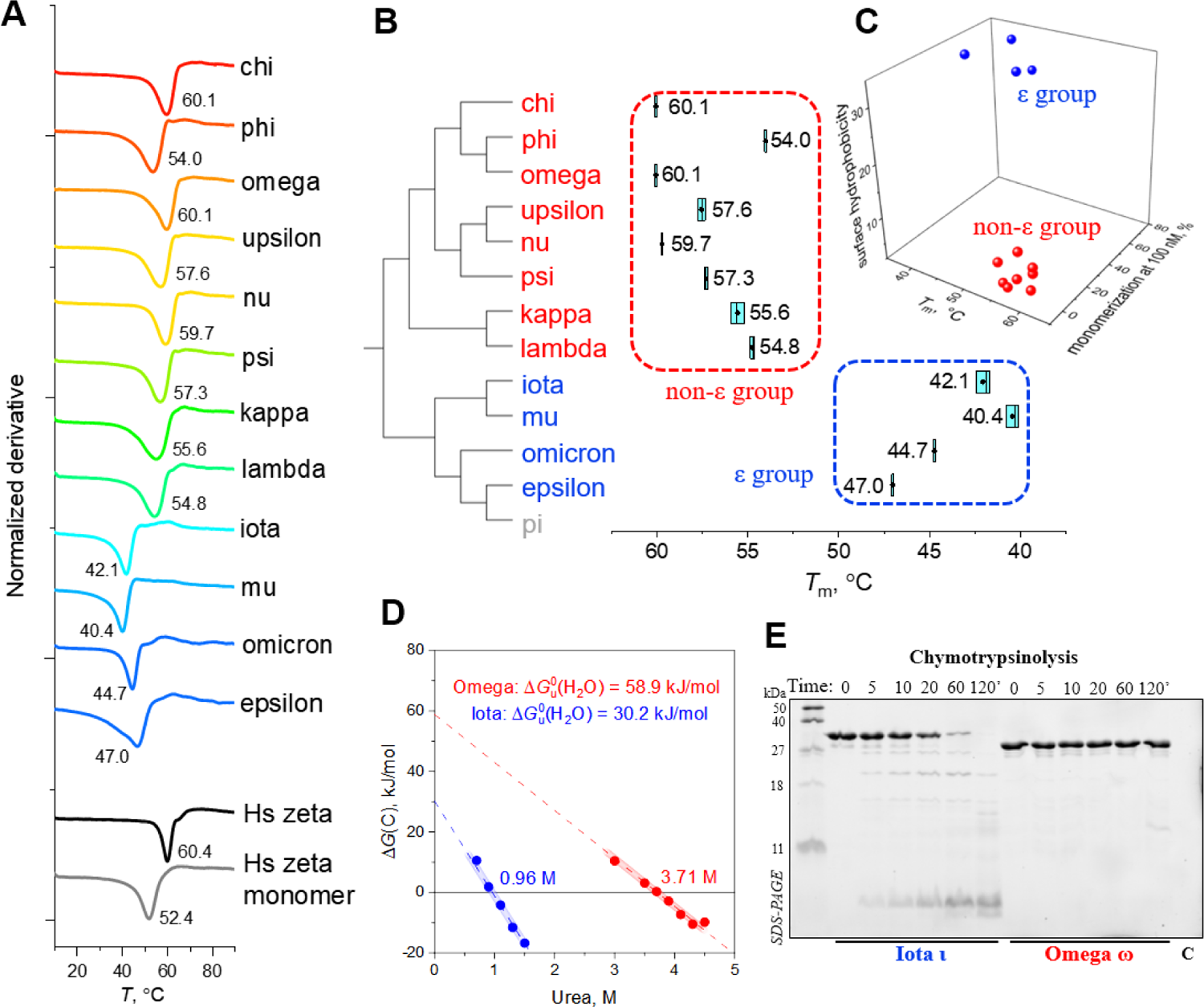
Epsilon-group *Arabidopsis* 14-3-3 isoforms display outstanding thermodynamic instability and undergo rapid proteolysis. **A**. Thermofluor-based stability profiling of the twelve Arabidopsis 14-3-3 isoforms compared with human 14-3-3 zeta wild-type (Hs zeta) and its monomeric mutant ζm58E (Hs zeta monomer). Final protein concentration in all cases was 2 μM (5 μM for epsilon). Heating from 10 to 90 °C (1 °C/min). Mean *T*_m_ values are shown (*n*=3). **B**. Thermal stability data for the *Arabidopsis* 14-3-3 isoforms robustly clusterize the epsilon and non-epsilon group isoforms recapitulating phylogenetic tree features. For convenience, a fragment of the phylogenetic tree is shown so that only its topology is preserved. The thermal stability data are presented as box plots. Line in the center indicates the median value, black dot shows the mean value, the box width corresponds to the mean +/- standard deviation (*n*=3). **C**. Clusterization of epsilon and non-epsilon isoforms in 3D using key biochemical signatures discovered in this work: thermal stability (*T*_m_), monomerization degree at 100 nM protein concentration, and surface hydrophobicity (bis-ANS fluorescence at 490 nm). **D**. Thermodynamics of iota and omega unfolding induced by urea, quantified according to the two-state model (Makhatadze, 1999). The urea concentration at which half of the protein is unfolded and values of the Gibbs energy of unfolding are indicated. **E**. Limited chymotrypsinolysis of *Arabidopsis* 14-3-3 iota and omega at a substrate:protease weight ratio of 200:1 at 27 °C. Protein concentration was 0.5 mg/ml.

To test whether the lower thermal stability of the epsilon-group isoforms is associated with the dissociation of the corresponding homodimers, we repeated thermal shift assay on 14-3-3 samples pre-crosslinked with BS3 under conditions yielding efficient formation of crosslinked dimers. Supplementary Fig. 9 shows that, while crosslinking induced moderate stabilization of omega (by ∼5 °C), *T*_m_ for iota increased from 42.1 to 56.6 °C and approached the *T*_m_ values of non-epsilon group homodimers of phi, lambda and kappa. Similar crosslinking-induced stabilization was observed for omicron, mu and epsilon, which confirmed the hypothesis that they display a dissociation mechanism of heat-induced denaturation (Supplementary Fig. 9). We did not reveal that ionic strength influences *T*_m_ values of selected isoforms any significantly (Supplementary Fig. 8), implying that electrostatic interactions play a limited role in overall stabilization of these 14-3-3 homodimers. The experimentally determined *T*_m_ values of *Arabidopsis* 14-3-3 proteins recapitulated clustering of epsilon and non-epsilon group isoforms on the phylogenetic relationships between all twelve *Arabidopsis* isoforms (Fig. 3B). Moreover, combining the three biochemical signatures, i.e. lowered thermal stability, increased monomerization propensity and enhanced surface hydrophobicity, we were able to graphically separate epsilon and non-epsilon phylogenetic groups on a 3D plot (Fig. 3C).

Given that the heat-induced denaturation of *Arabidopsis* 14-3-3s depended on dimer dissociation yet no substantial effect of ionic strength could be found, one could suspect the role of H-bonds in homodimer stabilization. To clarify this, we studied thermodynamics of 14-3-3 unfolding induced by urea, which revealed unfolding of iota occurring at dramatically lower urea concentration (0.96 M) compared to omega (3.71 M), with nearly twice as lower Gibbs energy of unfolding and much sharper denaturation curve (Fig. 3D and Supplementary Fig. 10). Of note, upon complete omega unfolding at ≥6 M urea concentration, we detected substantial protein refolding into dimers and monomers during native PAGE (Supplementary Fig. 10), which indicated the reversibility of urea-induced denaturation of these 14-3-3s and supported the applicability of equilibrium thermodynamics formalism.

The beginning of thermal transition of the epsilon-group 14-3-3 isoforms occurred at as low as 30-40 °C, i.e., rather close to the physiological temperatures experienced by plants (Antoun and Ouellet, 2013). In accord, we observed marked aggregation propensity of mu and iota compared with omega upon incubation of these isoforms at 37, 40 and 60 °C (Supplementary Fig. 10). Indeed, while omega remained intact for at least 2 h at 37 °C, 0.5 h at 40 °C and 1 min at 60 °C, mu started to aggregate already after 30 min incubation at 37°C. Likewise, iota started to aggregate after 5 min at 40 °C and completely aggregated after 1 min at 60 °C, whereas this high temperature caused significant aggregation of omega only after 30 min incubation (Supplementary Fig. 10).

The lowered thermodynamic stability of epsilon-group 14-3-3 isoforms suggested that they can undergo facile degradation. As a proxy, we subjected all isoforms to chymotrypsinolysis, which revealed that at least three out of four epsilon-group isoforms (except omicron) were almost completely degraded under conditions (2h 27 °C) when all non-epsilon isoforms remained intact (Fig. 3E and Supplementary Fig. 11).

### Structure-guided mutagenesis of 14-3-3 iota rescues the non-epsilon behavior

Careful structural comparison of 14-3-3 omega and iota and inspection of multiple sequence alignment of the *Arabidopsis* 14-3-3 isoforms allowed us to spot several positions that are occupied by functionally different residues in epsilon and non-epsilon groups (Fig. 4A-C). First, a hydrophobic core-stabilizing Phe residue found in all non-epsilon *Arabidopsis* 14-3-3 isoforms is replaced by Thr in iota and by other non-aromatic residues in other epsilon-group isoforms (Fig. 4A). Second, an interfacial His residue conserved in all non-epsilon group isoforms is replaced by Asn in epsilon group isoforms (Fig. 4B). Another, less apparent difference is found in the third α-helix of 14-3-3: while in all non-epsilon isoforms of *Arabidopsis* the corresponding position is occupied by a helix-promoting Ala, all epsilon group isoforms contain a helix-destabilizing Gly residue in this position (Fig. 4C). To investigate the effect of each of these replacements on the epsilon-type behavior of 14-3-3 iota, we prepared and analyzed three single mutants thereof, T32F, N84H and G54A.

**Fig. 4.**
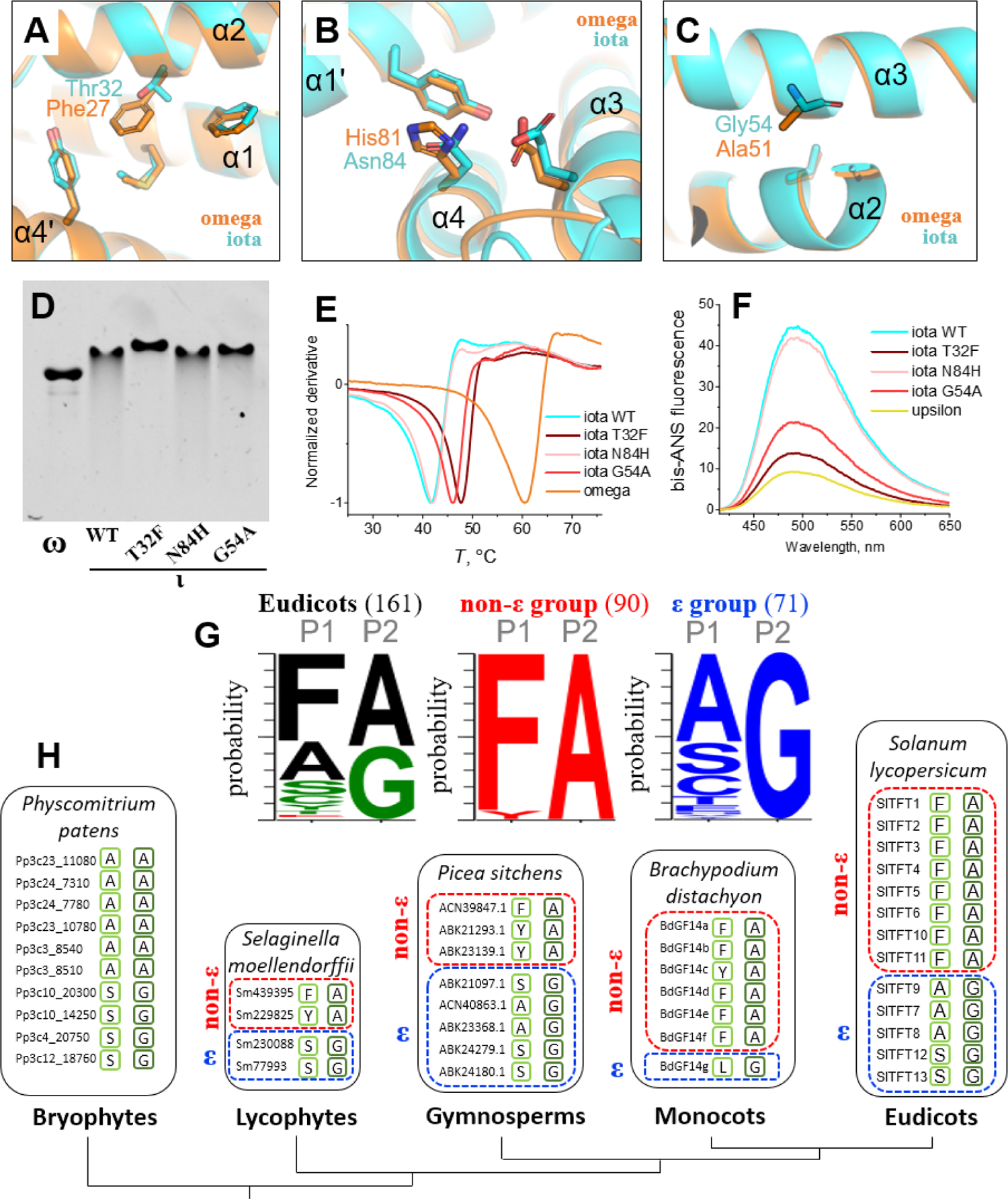
Decoding of biochemical signatures demarcating epsilon and non-epsilon phylogenetic groups of plant 14-3-3s. **A-C**. Structural comparison of 14-3-3 omega (PDB 8QTT) and iota (AlphaFold model) around the tentative demarcating positions. Alpha-helices are labeled. **D**. Native PAGE of omega, iota WT and single iota mutants T32F, N84H, G54A. **E**. Thermal stabilities of omega, iota WT and single iota mutants T32F, N84H, G54A determined by thermal shift assay at 1 °C/min heating rate from 10 to 90 °C. **F**. Surface hydrophobicity of upsilon, iota WT and single iota mutants T32F, N84H, G54A probed by bis-ANS. **G**. WebLogo (Crooks *et al*., 2004) diagrams showing the amino acids found in positions P1 (residue 27 in omega) and P2 (residue 51 in omega) in 161 isoforms of 14-3-3 from 13 sequenced Eudicot species (*A. thaliana, Glycine max, Nicotiana tabacum, Malus domestica, Solanum lycopersicum, Camellia sinensis, Manihot esculenta, Hevea brasiliensis, Theobroma cacao, Coffea arabica, Populus trichocarpa, Vitis vinifera, Medicago truncatula*) (Mikhaylova *et al*., 2021; Jia *et al*., 2022; Zhang *et al*., 2022; Zuo *et al*., 2021; Chang *et al*., 2020; Tian *et al*., 2015). Note the wide co-occurrence of the Phe (position P1) and Ala (position P2) in non-epsilon groups, suggesting usefulness of a FA criterion for the epsilon/non-epsilon demarcation. The WebLogo diagrams after division into epsilon and non-epsilon groups are also presented. **H**. The applicability of the FA criterion for the prediction of properties of 14-3-3 isoforms from various plants from Eudicots to Lycophytes, with exemplary organism 14-3-3 sets shown for every plant group. Positions P1 and P2 are outlined by bright and dark green, respectively. For Bryophytes, classification into epsilon and non-epsilon groups is still uncertain due to the absence of aromatic residues in P1.

As we hoped, the T32F and G54A substitutions completely (T32F) or partly (G54A) eliminated the monomerization of iota on native PAGE (Fig. 4D), increased its *T*_m_ value by 6.0 (T32F) or 4.3 °C (G54A) (Fig. 4E) and substantially reduced its surface hydrophobicity probed by bis-ANS (Fig. 4F). On the contrary, the effect of the N84H mutation was negligible, indicating that variation at this position is dispensable for 14-3-3 dimer stability.

### Key positions demarcate the epsilon versus non-epsilon properties across plant groups

Mutagenesis data indicated the importance of position P1 (occupied by Phe27 in omega) and position P2 (Ala51 of omega) for the epsilon/non-epsilon demarcation. We have analyzed over 150 sequences of 14-3-3 isoforms from more than 10 Eudicot species and revealed that for all cases, the known non-epsilon isoforms do contain Phe (or, rarely Tyr) in position P1 and Ala in position 2, which can be formulated as an ‘FA criterion’ (Fig. 4G). Epsilon-group isoforms always contain Gly in position P2 and a non-aromatic residue in position P1. We then asked if such descriptor could be used to determine 14-3-3 isoform affiliation with either epsilon or non-epsilon phylogenetic/functional groups in plant species beyond Eudicots. By choosing several representative organisms of Eudicots, Monocots, Gymnosperms, Lycophytes and Bryophytes we analyzed their 14-3-3 isoform sets using the FA criterion and found that it works in most cases from Eudicots to as far as Lycophytes (Fig. 4H). Very importantly, the FA criterion worked in the case of *Selaginella* (Lycophytes), which has four isoforms falling into epsilon (two) and non-epsilon (two) groups. Interestingly, based solely on phylogenetic analysis Cao et al reported that *Selaginella* has only four epsilon-group isoforms (Cao and Yan, 2016), whereas Zhang et al described *Selaginella* 14-3-3 isoforms as represented by both epsilon and non-epsilon types (Zhang *et al*., 2022). In the case of Bryophytes, we observed either an Ala or a Gly residue in position P2, but a mixture of non-aromatic residues in position P1, somewhat violating the FA rule. Properties of these proteins remain to be investigated, and it is likely that the demarcation of epsilon/non-epsilon groups obeys molecular mechanisms alternative to the FA criterion, which nonetheless performs well on most of the analyzed species/isoform sets.

## Discussion

Here we describe the discovery of the set of biochemical signatures robustly demarcating epsilon and non-epsilon groups of *Arabidopsis* 14-3-3 isoforms including the pronounced homodimer instability, monomerization, increased surface hydrophobicity, proteolytic degradation propensity and aggregation of the former group. Data mining using Protein Abundance Database (PaxDB (Wang *et al*., 2015)), allowing us to estimate effective intracellular concentrations of the studied isoforms, confirmed that the observed phenomena are manifested at physiologically relevant, low-micromolar protein concentrations (Fig. 1D and Table 1). Therefore, our findings have far reaching consequences for understanding 14-3-3 protein biology in plant physiology.

**Table 1.**
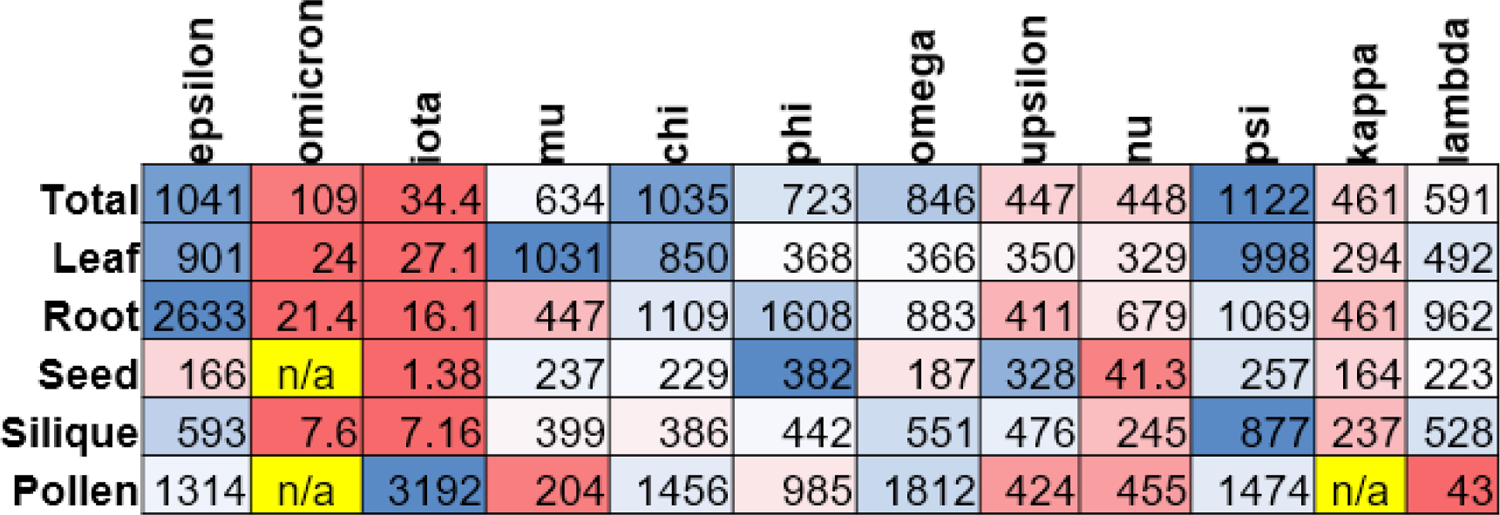
Organ-specific distribution of twelve 14-3-3 isoforms in *A. thaliana*. The numbers represent ppm values retrieved from PaxDB (Wang *et al*., 2015). n/a - data not available. 1 ppm equals 1 part per million of total protein molecules and can be converted into molar concentration assuming total concentration of protein molecules of ∼3 mM (Milo, 2013). “Total” corresponds to the abundance of specific isoforms in the whole organism. The rows are color coded according to the gradient from low (0%, red), to intermediate (50%, white), to high (100%, blue) using a conditional formatting tool in Microsoft Excel.

It is generally accepted that eukaryotic 14-3-3 isoforms are conservative dimers although an accurate comparative analysis has not been undertaken even for the well-studied model organisms. A quantitative measure of the affinity between subunits within a 14-3-3 dimer has been reported only for human 14-3-3 zeta (*K*_D_= 5 nM) (Trošanová *et al*., 2022). While dimerization is believed to be important for most of 14-3-3 functions (Y., Shen *et al*., 2003; Messaritou *et al*., 2010), several works suggested monomer-specific functions (Zhou *et al*., 2003; Sluchanko and Gusev, 2017; Sluchanko *et al*., 2014; Zhu *et al*., 2023) and their regulation via alteration of the dimer versus monomer status. Nonetheless, this status remains unexplored for most of the 14-3-3 members, including particularly numerous plant 14-3-3s (Mikhaylova *et al*., 2021; Zhang *et al*., 2022).

The choice of *A. thaliana* for this work was based on the fact that it is the well-characterized plant, with rich high-throughput data available, including the data on the abundance of each isoform in the whole plant and in distinct organs (see Table 1) (Wilson *et al*., 2016; Wang *et al*., 2015). According to PaxDB (Wang *et al*., 2015), *A. thaliana* 14-3-3 isoforms are nonuniformly expressed in the plant organs, and there is a great variation in the total 14-3-3 abundance (Fig. 1D and Table 1). Estimating the overall intracellular concentration of proteins as 3 mM (Milo, 2013), one can convert the available abundance values in ppm into micromolar concentration of the protein of interest (Wang *et al*., 2015; Gogl *et al*., 2021). Such estimation reveals that the expression level of *Arabidopsis* 14-3-3 isoforms can differ by 34 times, from only 100 nM for iota to 3.4 μM for psi (Table 1). There is no strict correlation in the expression levels for the epsilon and non-epsilon groups, as the most abundant isoforms are found in either group (epsilon and chi/psi, respectively). However, the overall least abundant are epsilon-group isoforms (iota and omicron). Of note, pi is the least abundant 14-3-3 isoform with a very low expression level restricted to anthers and chalazal cyst of the seed (Keicher *et al*., 2017). Our attempts to purify 14-3-3 pi have so far been unsuccessful, likely because of the presence of the largest number of cysteines among *Arabidopsis* 14-3-3 isoforms (5 compared to 2 in others) and its extreme instability.

Surprisingly, the pattern of the *T*_m_ values for twelve *Arabidopsis* isoforms correlate with their phylogenetic tree, reflecting not only the major division into epsilon and non-epsilon groups, but also relationships within the groups (Fig. 1D). For instance, among epsilon-group isoforms, higher thermostable are omicron and epsilon, together forming epsilon-subgroup, whereas thermal stabilities within subgroups psi (57.3-59.7 °C) and kappa (54.8-55.6 °C) of the non-epsilon group are also very close to each other and different between. In the omega subgroup, one exception is exhibited by phi, which is much less stable (54 °C) than the other two isoforms of the same subgroup (60.1 °C each). Variations of quantifiable properties within the phylogenetic 14-3-3 subgroups emphasize the importance of systematic comparative analyses of all members of the 14-3-3 family and highlights risks associated with the analysis of voluntarily picked individual representatives, as such widely practiced approach (Lambeck *et al*., 2010; Rosenquist *et al*., 2000; Sullivan *et al*., 2009; Pallucca *et al*., 2014) can produce borderline results blurring functional categories. For instance, the *T*_m_ difference between the highest stable epsilon-group isoform (epsilon, 47 °C) and the lowest stable non-epsilon group isoform (lambda, 54.8 °C) is only ∼7.8 °C, which is much less contrast than ∼20 °C difference between other representatives of these groups, i.e. mu and chi/omega. Likewise, the smearing electrophoretic pattern associated with the monomerization of omicron and its proteolytic pattern are least pronounced among the four epsilon-group isoforms and could have been overlooked if compared with any non-epsilon isoform alone. Omicron also shows the least hydrophobicity among the epsilon-group isoforms, which is only ∼2.7 times larger than that of the highest hydrophobic non-epsilon isoform, upsilon. It matches the greatest difference within the non-epsilon group (upsilon vs psi; ∼3 times), and is much less contrasting than the ∼10-times difference between epsilon-group iota and non-epsilon group psi. Nevertheless, systematic cross-family comparison enabled unequivocal clusterization of omicron with the rest epsilon-group isoforms (Fig. 3C). As mentioned, the largest *T*_m_ difference between the epsilon and non-epsilon group isoforms covers almost 20 °C (mu vs omega/chi). The occurrence of the thermal transition at relatively low temperatures (*T*_m_ of 40-42 °C) indicate a particular physiological relevance of the observed instability ‘encoded’ in the epsilon-group, as these isoforms tend to aggregate already at 35-37 °C, which is close to ambient temperatures (Antoun and Ouellet, 2013). Diversification of thermal stabilities of 14-3-3 isoforms may be exploited by plants for isoform- and tissue-specific tasks, especially given that distinct plant organs can experience substantially different temperatures (Fu *et al*., 2016; Yan *et al*., 2008). Of note, a recent study showed that transcription of tomato 14-3-3 isoforms is up-regulated upon chilling (4°C) and, conversely, downregulated by continuous heat stress (42 °C), whereas overexpression of one of the non-epsilon isoforms (SlTFT6) improves heat tolerance (Liang *et al*., 2023).

Deep monomerization of iota and omicron is intriguing given that they are the least abundant 14-3-3 isoforms in *Arabidopsis*, with average intracellular concentrations of 100-300 nM (Fig. 1D). Moreover, concentration of these isoforms is even lower in silique, seed, root and leaf (Table 1), which makes the consequences of the discovered dimer instability of these isoforms even more relevant. At the same time, iota is highly abundant in pollen (3192 ppm, ∼10 μM (Wang *et al*., 2015)) and is the highest abundant there among other isoforms (Table 1). This points to tissue-specific functions of 14-3-3 iota in the pollen, especially in the light of the recent data on overrepresentation of phosphoproteins containing 14-3-3-binding motifs in the pollen, and therefore suggests important functions of 14-3-3 proteins in pollination (Mayank *et al*., 2012). Interestingly, the reported abundance of iota is minimal in the *Arabidopsis* seed (1.38 ppm, ∼4 nM), which is ∼280 times lower than the abundance of phi in the same organ (Table 1). 14-3-3 proteins change expression levels under the conditions of the biotic and abiotic stress (Zhang *et al*., 2022). This strongly indicates that the level of expression of a given isoform is one of the key regulatory means, and variations in the range of concentrations where isoforms display instability of their dimers can be most efficient. Our data strongly suggest that the half-life of epsilon-group 14-3-3 isoforms can be limited by their faster degradation associated with lower thermodynamic stability.

Our work culminated in using site-directed mutagenesis to decode that biochemical peculiarities discovered here for the epsilon-group 14-3-3 isoforms (monomerization and aggregation propensity, thermodynamic instability, higher surface hydrophobicity and higher susceptibility to proteolytic degradation) are mainly dictated by two amino acid positions, Thr32 (P1) and Gly54 (P2) (iota numbering), whose mutation in iota largely restores the non-epsilon-like behavior. Structural data nicely rationalize these effects and suggest that a combination of residues found in P1 and P2 can robustly predict affiliation of a given 14-3-3 isoform to either phylogenetic group, which is especially useful for poorly described plant isoforms. The widely used traditional phylogenetic analysis based on entire 14-3-3 sequences (Liang *et al*., 2023; Zuo *et al*., 2021; Jia *et al*., 2022; Chang *et al*., 2020; Tian *et al*., 2015; Zhang *et al*., 2022; Cao and Yan, 2016; Ren *et al*., 2023) is rather weak on its own, particularly for evolutionarily distant species, for which it provides controversial and unreliable results. For example, Lycophyte *Selaginella* has been reported to contain either only epsilon-group or both epsilon and non-epsilon group isoforms (Cao and Yan, 2016; Zhang *et al*., 2022), whereas the combination of residues occupying P1 and P2 positions robustly demarcate two epsilon and two non-epsilon isoforms in this plant. In Bryophytes, the mixture of amino acids in P1 and P2 positions is observed, which may suggest the existence of alternative mechanisms for 14-3-3 dimer stabilization in these plants, which warrants further investigations. Notwithstanding, our approach based on P1 and P2 positions facilitates demarcation of previously unaddressed or controversially divided functional 14-3-3 types in Gymnosperms and Monocots, while also showing excellent performance on Eudicot 14-3-3 isoforms, fully recapitulating their phylogenetic categorization into epsilon and non-epsilon groups. Thus, our study provides a framework for the multiway comparisons, analyses and predictions that will hopefully reinvigorate future research of plant 14-3-3s.

## Materials and Methods

### Materials

All regular chemicals were of the highest purity and quality available. Isopropyl-β-thiogalactoside (IPTG) was from Thermo Scientific (#R0392), suberic acid bis(3-sulfo-N-hydroxysuccinimide ester) sodium salt (BS3) was from Sigma (#S5799), 4,4’-bis(phenylamino)-[1,1’-binaphthalene]-5,5’-disulfonic acid (bis-ANS) was from MP Biomedicals (#150097), thrombin was from Tehnologia-standart (#017).

### Molecular cloning

Molecular constructs encoding *A. thaliana* 14-3-3 isoforms nu and upsilon were obtained by reverse transcription from *A. thaliana* cell lysates. Cell culture material was provided by the Laboratory of plant cell culture biology of Institute of plant physiology of the Russian Academy of Sciences. Suspension cell cultures of *A. thaliana* (L.) Heynh. wild type (Col-0) were generated by Drs. A.V. Nosov and A.A. Fomenkov and deposited at the All-Russia Collection of Cultivated Cells of Higher Plants as NFC-0. Cells were cultured in 50 mL of Schenk and Hildebrandt medium (Schenk and Hildebrandt, 1972 (Schenk and Hildebrandt, 1972)) supplemented with 3% sucrose, 1 mg/L 2,4-Dichlorophenoxyacetic acid (2,4-D) (Sigma, St. Louis, Missouri, USA) and 0.1 mg/L kinetin (Sigma). Cells were grown for 10 d under constant agitation (110 rpm) at 26 °C and 70% humidity (Fomenkov *et al*., 2014). The cells were separated from the culture medium by filtration through a paper filter, frozen in liquid nitrogen and then ground to fine powder with mortar and pestle (both were pre-cooled with liquid nitrogen). The powder was stored at −80 °C. Total RNA was isolated with Extract RNA reagent (Evrogen) according to the manufacturer’s protocol. The RNA quality was controlled by agarose gel electrophoresis. Total RNA obtained was subjected to reverse transcription using Magnus reverse transcriptase (Evrogen) with oligodT primers (Evrogen) according to the manufacturer’s protocol. The resulting total cDNA was subjected to PCR with primers Upn_for, Upn_rev for amplification of the upsilon coding sequence and Nu_for, Nu_rev for the nu coding sequence (all primer sequences are found in Supplementary Table 1). Forward primers contained the *Nde*I restriction site and reverse primers contained the *Xho*I restriction site. The PCR was performed with *Pfu* DNA polymerase (Sibenzyme). Specific PCR products were cleaned-up from agarose gels and digested with *Nde*I and *Xho*I enzymes (Sibenzyme). The resulting DNA molecules were ligated into a pET28a vector (kanamycin resistance) with Instant Sticky-end Ligase Master Mix (NEB) and transformed for amplification in DH5α *E. coli* cells. Insertion of a specific gene was verified by Sanger sequencing using T7_forward and T7_reverse primers of the purified plasmids (DNA sequencing was performed at the Shared-Access Equipment Centre “Industrial Biotechnology’’). The pET28a based molecular constructs encoding *A. thaliana* 14-3-3 isoforms omega, mu, epsilon, iota, and pi were obtained by gene synthesis (Synbio). Molecular constructs encoding *A. thaliana* 14-3-3 isoforms chi, psi, phi, lambda, kappa and omicron were kindly provided by Prof. Robert J. Ferl.

Sanger sequencing using T7_forward and T7_reverse primers revealed that 14-3-3 chi had six N-terminal amino acids altered compared to the canonical Uniprot variant. Therefore, we constructed a new specific forward primer to add the correct N-terminal sequence, re-amplified the chi coding region using Chi_for, Chi_rev primers (harboring *Nde*I-*Xho*I sites) and inserted the endonuclease-digested PCR product into pET28a vector. The correctness of the insert was verified by Sanger sequencing. The constructs were transformed into chemically competent *E. coli* C41(DE3) cells (psi, phi, lambda, kappa and omicron) or BL21(DE3) (all other isoforms) cells for protein expression.

Experimental procedures with 14-3-3 pi required additional manipulations to obtain a maltose-binding protein (MBP)-tagged fusion. The 14-3-3 pi coding sequence surrounded by restriction sites was obtained from pET28a construction via *Nde*I-*Xho*I digestion of the pET28a-14-3-3 pi plasmid. The construct was cloned into a pETX-MBP vector (kanamycin resistance, 3C cleavage site after MBP). The plasmids were transformed into chemically competent *E. coli* BL21(DE3) cells for protein expression.

14-3-3 iota mutants were obtained by the megaprimer PCR method. At the first step pET28a-14-3-3 iota construction was amplified with T7_for and M1_rev or M2_rev or M3_rev partially complementary primers to obtain megaprimer oligonucleotides (M1, M2, M3 megaprimers respectively; M1 for T32F site-directed mutagenesis, M2 for N84H and M3 for G54A). At the next step, the purified M1, M2, M3 megaprimers and T7_rev primers were introduced in the PCR with pET28a-14-3-3 iota as a matrix. The resulting amplified product was digested with *Nde*I/*Xho*I and cloned into pET28a. The presence of the specific mutation was confirmed by Sanger sequencing. Resulting plasmids were transformed into chemically competent *E. coli* BL21(DE3) cells for protein expression.

### Protein expression and purification

For His_6_-tagged 14-3-3 chi, omega, psi, phi, upsilon, lambda, nu, kappa, epsilon, omicron, iota (and mutants of the latter), protein expression was initiated by the addition of isopropyl-β-thiogalactoside (IPTG) to a final concentration of 0.4 mM and continued for 16 h at 28 °C. For 14-3-3 mu we found that expression initiated at 0.1 mM IPTG and continued for 16 h at 18 °C led to a substantially higher protein yield in the soluble fraction. 14-3-3 iota single a.a. mutants expression was initiated with 0.1 mM IPTG and continued for 16 h at 18 °C. The expressed His_6_-tagged proteins were purified in 3 steps: 1) immobilized metal-affinity chromatography (IMAC) with subsequent overnight thrombin-assisted cleavage of the His_6_-tag, 2) subtractive IMAC and 3) gel-filtration using Superdex 200 26/600 (GE Healthcare). Specially for SEC-MALS analysis, we took aliquots of each isoform before preparative SEC to capture all oligomeric forms potentially present. For other methods we used 14-3-3 samples polished by preparative SEC. Purified protein was concentrated to 20-25 mg/ml and stored in aliquots at −80 °C.

Expression of His_6_-tagged 14-3-3 pi in *E. coli* BL21(DE3) cells was initiated by adding IPTG up to the final concentration of 0.05 mM and continued for 16 h at 18 °C. However, the overexpressed protein was present almost entirely in the insoluble fraction. Subtractive IMAC yielded a low amount of protein which was substantially contaminated with a 60 kDa protein likely corresponding to GroEL.

Expression of MBP-14-3-3 pi protein was induced by 0.05 mM IPTG and continued for 16 h at 18 °C. Almost all obtained fused protein was soluble and was purified via MBP-trap column. The resulting eluate was subjected to 3C proteolysis. However, the protein was digested only partly in spite of the fact that the protease aliquote was added twice to ensure the efficient proteolysis. Analytical SEC showed that the protein was present in aggregates. The identity of the twelve purified plant 14-3-3 isoforms and the intactness of the protein termini were verified by MALDI-TOF mass-spectrometry on a Bruker UltraFlexTreme mass spectrometer (Supplementary Fig. 1).

Untagged full-length human 14-3-3ζ and its monomeric phosphomimetic mutant ζm58E were expressed in *E. coli* BL21(DE3) cells by IPTG induction (final concentration of 1 mM) and purified by ammonium sulfate fractionation, anion-exchange chromatography and SEC as described earlier (Sluchanko and Uversky, 2015; Sluchanko *et al*., 2008). His-tagged full-length human 14-3-3ε was obtained in previous work (Tugaeva *et al*., 2020). Lambda phosphatase was obtained as a recombinant protein with the N-terminal His_6_-tag cleavable by human rhinovirus 3C protease in previous work (Kapitonova *et al*., 2024). The dephosphorylation reaction (20 µl) lasted 5 h at 20 °C and contained 1 mg/ml of purified plant 14-3-3 samples or pre-phosphorylated HSPB6 in 20 mM Tris-HCl pH 7.8 with 150 mM NaCl, 1.9 mM MnCl_2_, 5 mM dithiothreitol (DTT) and 1/10 diluted lambda phosphatase. Protein concentrations were determined on a NP80 nanophotometer (Implen, Munich, Germany) at 280 nm using the extinction coefficients listed in Supplementary Table 2.

### Size-exclusion chromatography

Dissociation of 14-3-3 homodimers upon the decrease of protein concentration was followed by loading serial dilutions prepared on SEC buffer (20 mM Tris-HCl, pH 7.5, 150 mM NaCl, 5 mM β-mercaptoethanol (βME), 3 mM NaN_3_) on a Superdex 200 Increase 5/150 column (GE Healthcare) equilibrated by the same buffer and run at a 0.45 ml/min flow rate. The elution profiles were followed by intrinsic fluorescence at 340 nm excited at 290 nm using a 1260 FLD spectra detector (G7121B, Agilent) and then normalized to 0,1.

To estimate the Stokes radii (*R*_S_) for the protein peaks we plotted the calibration curve (-log *K*_av_)^0.5^ against *R*_S_ in nm based on hydrodynamic radii of BSA trimer (5.4 nm), BSA dimer (4.6 nm), BSA monomer (3.5 nm). *K*_av_ values for BSA and 14-3-3 peaks were determined as (*V*_e_-*V*_0_)/(*V*_t_-*V*_0_), where *V*_e_ is elution volume at peak apex (in ml), *V*_t_ = 24 ml and *V*_0_ = 8 ml. In addition, we estimated apparent *M*_w_ for the 14-3-3 peaks using BSA dimer (132 kDa), BSA monomer (66 kDa) and α-lactalbumin (15 kDa). To estimate the extent of the dimer-to-monomer transition for each 14-3-3 isoform analyzed, we first considered the apparent *M*_w_ at 100 µM protein concentration as *M*_w_ for a pure dimer, and a twice lower value as *M*_w_ for a pure monomer (assuming a similar protein shape). Then, the elution volumes of each 14-3-3 isoform at a given concentration were converted into the apparent *M*_w_ values and represented at the fraction of the dimer-to-monomer transition.

### SEC-MALS

Absolute molecular masses of the twelve 14-3-3 isoforms were determined independently of a protein shape and possible nonspecific interactions with the chromatographic resin, by SEC coupled with multi-angle light scattering (SEC-MALS). To this end, each 14-3-3 sample purified to homogeneity but before preparative SEC was prepared on SEC buffer (65 µl; ∼3 mg/ml or 100 μM per monomer), clarified by centrifugation for 5 min at 21,400 g at 4 °C, pre-incubated for at least 30 min at 6 °C within the Vialsampler (G7129A, Agilent), and then loaded on a Superdex 200 Increase 10/300 column (Cytiva) pre-equilibrated with the same buffer. The runs at 0.8 ml/min flow rate were performed on an Agilent 1260 Infinity II chromatography system equipped with a 1260 Infinity II WR diode-array detector (G7115A, Agilent), a miniDAWN detector (Wyatt Technology) and a refractometric detector (G7162A, Agilent, operated at 35 °C), connected by the thinnest PEEK tubings available, in the indicated order. The 0.1 µm post-column filter (Malvern) was used to reduce the noise detected by the miniDAWN detector. The elution profiles were followed by absorbance at 280 nm and changes of the refractive index, as well as static light scattering at three angles. All these data were collected and processed in Astra 8.0 software (Wyatt Technology) using the refractometric detector as a concentration source (dn/dc was taken equal to 0.185 for each protein) and 250,000 attenuation of the output RI signal. Normalization of the static light scattering signals at different angles detected by miniDAWN was done using a pre-run of a BSA standard (Wyatt). The polydispersity index (*M*_w_/*M*_n_) was nearly equal to 1.000 for all samples characterizing monodisperse particles and was not presented.

### Native gel electrophoresis

Native polyacrylamide gel electrophoresis was performed in the homogenous Tris-Glycine buffer system according to the Shaub and Perry method (Schaub and Perry, 1969) with modifications. Anode, cathode and the gel buffer were 80 mM glycine, titrated with Tris to pH 8.6. The gel consisted of two layers: the sample gel (5% acrylamide (AA)) and the resolving gel (15% AA or 15-25% linear gradient AA). The gels were customized in BioRad Mini-PROTEAN system, 1 mm thick, 15 wells. Samples were dissolved in SEC buffer to concentration 0.33 mg/ml, then mixed with 4x sample buffer and loaded into the wells in quantities of 0.4 μg of each protein sample. Electrophoresis was run for 2 h at room temperature at constant current conditions (6 mA per one gel), voltage was limited to 350 V. The protein bands were visualized in Gel Doc EZ System (BioRad) by Stain-Free ® technology using trichloroethanol (Appscience) put into the gel solution prior to polymerization.

### Chemical crosslinking

Protein samples were dissolved in the crosslinking buffer (20 mM HEPES-NaOH, 150 mM NaCl, pH 7.5) to obtain the final protein concentration of 13.3 μM. Small aliquots of stock solution of bis(sulfosuccinimidyl) suberate (BS3, Sigma) on the same buffer were added to the protein samples to obtain 0.7 mM BS3 concentration (∼50-fold molar excess over a protein). The reactions were carried out in 50 μL at 20 °C for 40 min. Then the reaction was quenched with addition of 1 M Tris-HCl pH 8.0, to reach 50 mM Tris concentration in the samples. Crosslinked proteins were subjected to native and SDS gel-electrophoresis and analyzed by thermal shift assay.

### Probing surface hydrophobicity of 14-3-3 proteins by bis-ANS

Surface hydrophobicity of 14-3-3 proteins (final concentration 2 μM) was analyzed by adding fluorescent environmentally sensitive probe bis-ANS (final concentration 10 μM) and recording steady-state fluorescence spectra in the range of 415-650 nm (excitation at 385 ± 20 nm; emission slit 10 nm). All measurements were performed at 25 °C on a Clariostar plus microplate reader (BMG Labtech, Offenburg, Germany) in 384-well plates (Black Nunc™ 384-Shallow Well Standard Height Polypropylene Storage Microplates, Thermofisher scientific, cat. no. 267460). Each 14-3-3 sample was mixed with a small aliquot of the stock solution of bis-ANS (∼300 μM in water) and split into two technical 50-μl aliquots. Then the plates were centrifuged for 3 min at 25 °C at 150 g to remove air bubbles, and the fluorescence spectra were recorded with a gain of 1,500. bis-ANS concentration in the stock solution was determined spectrophotometrically at 385 nm using the molar extinction coefficient equal to 16,700 M^−1^ cm^−1^. The protein concentration was equalized by absorbance at 280 nm and verified by SDS-PAGE of the aliquots of the reaction mixture with bis-ANS.

### Thermal shift assay

Protein samples (40 μl, final protein concentration 2 μM, unless the otherwise is stated) were mixed with ProteOrange (Lumiprobe; final concentration 5X) in 20 mM HEPES-NaOH buffer pH 7.5 containing 150 mM NaCl (unless NaCl concentration was varied to specifically study the effect of ionic strength), and the melting curves were monitored by ProteOrange fluorescence (excitation and emission filters 470 ± 15 nm and 586 ± 10 nm, respectively) using a QuantStudio 5 PCR machine (Applied Biosystems, ThermoFisher Scientific) at a heating rate of 1 °C/min. Before the run the samples were preconditioned to 10 °C for 10 min within the machine as part of the heating program. The first derivative of the melting curves was obtained to facilitate the analysis of thermal shifts. All samples were run in triplicates.

Additionally, we analyzed the effect of chemical crosslinking of 14-3-3 samples (1 μM) by BS3 on the denaturation profiles. Control samples included human 14-3-3ζ WT, human 14-3-3ε WT and the monomeric mutant form of 14-3-3ζ, ζm58E (Sluchanko and Uversky, 2015). Thermal shift characteristics obtained using ProteOrange dye were verified by monitoring intrinsic protein tryptophan fluorescence upon gradual heating in Cary Eclipse fluorimeter (Varian, Australia, Inc.). Protein samples (500 μl, proteins dissolved in 20 mM HEPES-NaOH buffer pH 7.5 containing 150 mM NaCl to final protein concentration 5 μM) were put into cuvettes, preincubated at 10 °C for 10 min and heated to 75 °C at a constant 1 °C/min heating rate. Fluorescence was excited at 297 nm and monitored at 340 nm (slits width 5 nm, detector voltage 700 V). Emission wavelength was selected according to tryptophan fluorescence spectra maximum; the spectra were obtained prior to the thermal shift analysis.

### Probing thermodynamic stability of 14-3-3 using urea denaturation

14-3-3 omega and iota were subjected to urea-induced unfolding by incubating protein samples (final protein concentration of 0.5 mg/ml) in the presence of various concentrations of urea (0-6 M) added as aliquots of the 10 M stock solution. After 5 h at 25 °C the samples were subjected to gel electrophoresis in 15 % PAAG run under otherwise non-denaturing conditions. The protein bands were visualized in Gel Doc EZ System (BioRad) by Stain-Free ® technology using trichloroethanol (Appscience) put into the gel solution prior to polymerization. Densitometric analysis in ImageLab 6.0 (Bio-Rad) yielded intensities of the native protein bands as a function of urea concentration. The sigmoidal curves obtained were fitted by the Boltzmann equation, to reveal the transition region. Then, the ratio of the fraction of unfolded protein to fraction of the folded protein at each urea concentration in the transition region, i.e. the equilibrium constant *K*_eq_, was used to determine Gibbs energy of the unfolding, Δ*G*_u_(C) = –RT ln *K*_eq_(C), by linear extrapolation to zero urea concentration from the linear region around the denaturation transition (Makhatadze, 1999). The reversibility of unfolding was assumed from the observation that at higher urea concentrations we observed substantial protein refolding in the gel, as a result of protein separation from urea in the course of native PAGE.

### Limited chymotrypsinolysis

To select suitable conditions of limited proteolysis, we first subjected 14-3-3 omega and iota (0.5 mg/ml) to chymotrypsinolysis at a substrate:protease weight ratio of 200:1 at 27 °C and stopped the reaction at selected time points by adding SDS sample buffer containing PMSF. The products of the reaction were separated by SDS-PAGE in 17% PAAG. Then, each of twelve Arabidopsis 14-3-3 isoforms (0.5 mg/ml) were subjected to the same treatment and the results were analyzed after 1 h and 2 h at 27 °C by SDS-PAGE in 17% PAAG.

## Data availability statement

All data associated with the study are available from the corresponding author upon reasonable request.

## Acknowledgements

The authors thank Prof. Robert J. Ferl for kindly providing pET15 plasmids encoding 14-3-3 chi, phi, kappa, lambda, psi and omicron from *A. thaliana* and Drs. A. A. Fomenkov and A. V. Nosov for providing *A. thaliana* cell culture materials. Cultivation of plant cell suspensions was performed using the equipment of the large-scale research facilities “Experimental biotechnological facility” and “All-Russian Collection of cell cultures of higher plants” of the Institute of Plant Physiology. The study was partly supported by the Ministry of Science and Higher Education of the Russian Federation (075-15-2021-1354). Sanger DNA sequencing, SEC-MALS and MALDI measurements were done at the Shared-Access Equipment Centre “Industrial Biotechnology’’ of the Federal Research Center “Fundamentals of Biotechnology” of the Russian Academy of Sciences.

## Author contributions

N.N.S. conceived and supervised studies, I.A.S. produced proteins and performed experiments, I.A.S. and N.N.S. analyzed data, N.N.S. wrote the paper with input from I.A.S.

## Supplementary Information

### Supplementary Tables

**Table S1.**
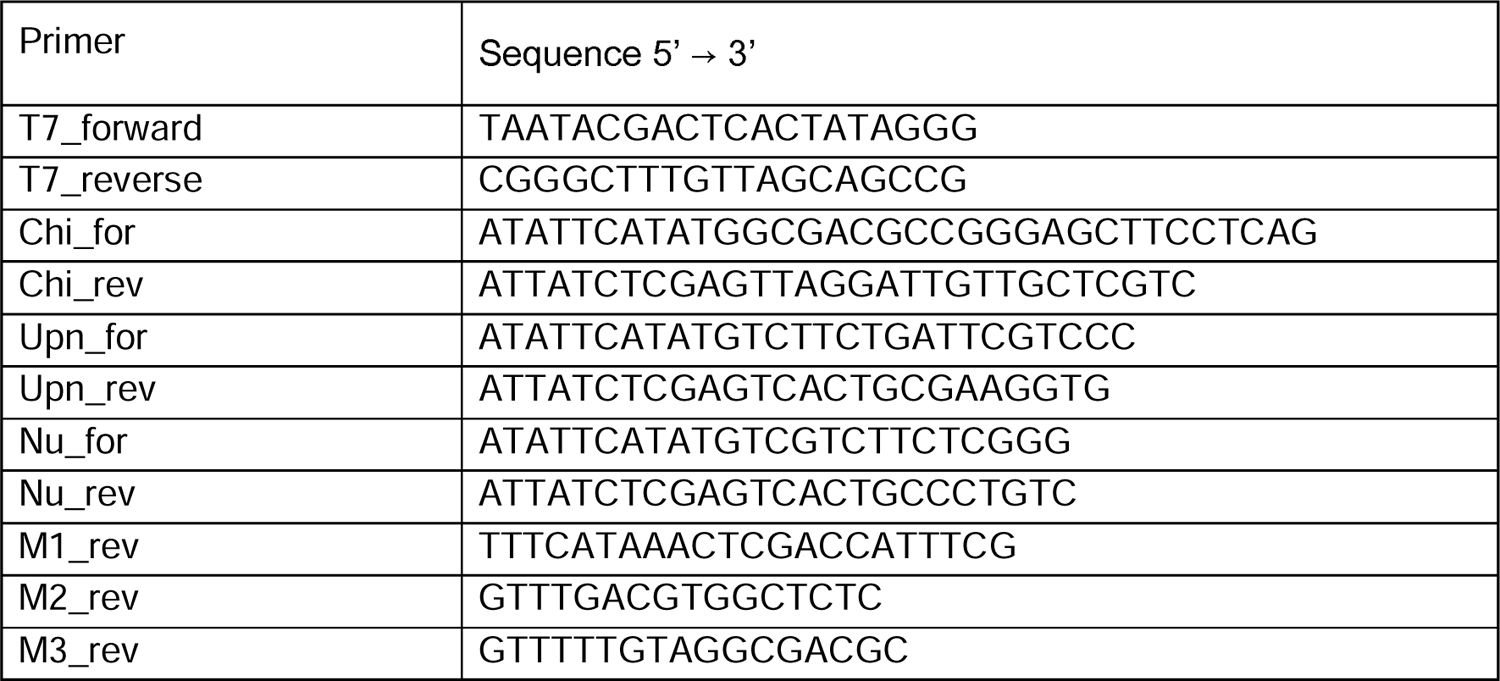
Primer sequences used in this study.

**Table S2.** Properties of the proteins studied.

| Protein | $M_w$, Da | pI | Molar ext. coef.<br>$M^{-1} \text{ cm}^{-1}$ | Ext. coef.<br>$(\text{mg/mL})^{-1} \text{ cm}^{-1}$ |
| --- | --- | --- | --- | --- |
| chi | 30213 | 4.73 | 27390 | 0.907 |
| omega | 29443 | 4.75 | 27390 | 0.930 |
| psi | 28887 | 4.77 | 27390 | 0.948 |
| phi | 30475 | 4.83 | 27390 | 0.899 |
| upsilon | 30463 | 4.77 | 27390 | 0.899 |
| lambda | 28257 | 4.82 | 28880 | 1.022 |
| nu | 30105 | 4.78 | 27390 | 0.910 |
| kappa | 28310 | 4.88 | 28880 | 1.020 |
| mu | 29801 | 4.89 | 27390 | 0.919 |
| epsilon | 29196 | 4.77 | 28880 | 0.989 |
| omicron | 29062 | 4.99 | 30370 | 1.045 |
| iota | 30826 | 4.88 | 30370 | 0.985 |
| iota T32F | 30872 | 4.88 | 30370 | 0.984 |
| iota N84H | 30849 | 4.92 | 30370 | 0.984 |
| iota G54A | 30840 | 4.88 | 30370 | 0.985 |
| human 14-3-3ζ | 27745 | 4.73 | 27390 | 0.990 |
| human His<sub>6</sub>-14-3-3ε | 32548 | 4.85 | 30370 | 0.930 |
| human 14-3-3ζm-S58E<br>(<sup>12</sup>LAE<sup>14</sup>→QQR) | 27886 | 4.78 | 27390 | 0.980 |

### Supplementary Figures

**Fig. S1.**
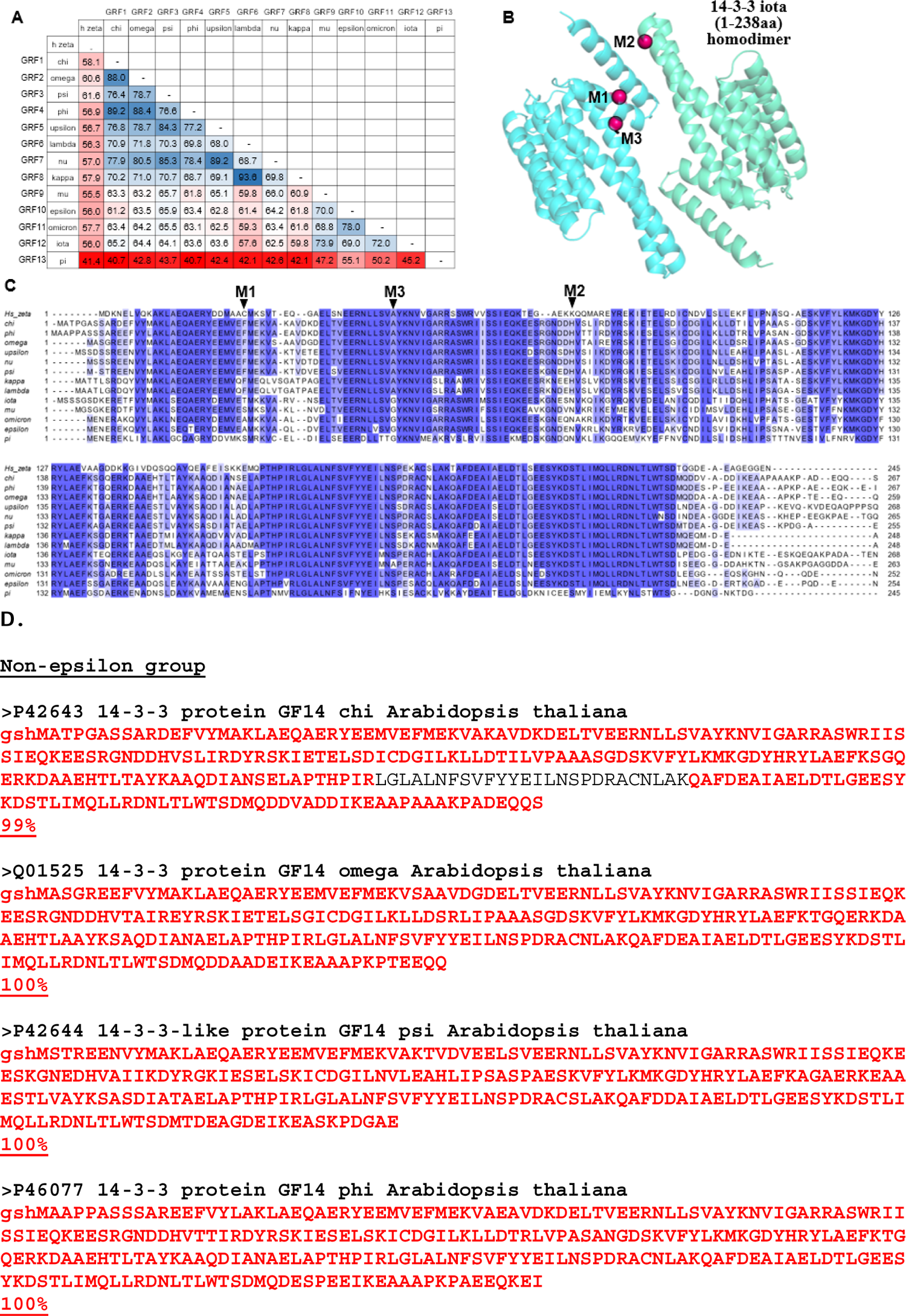
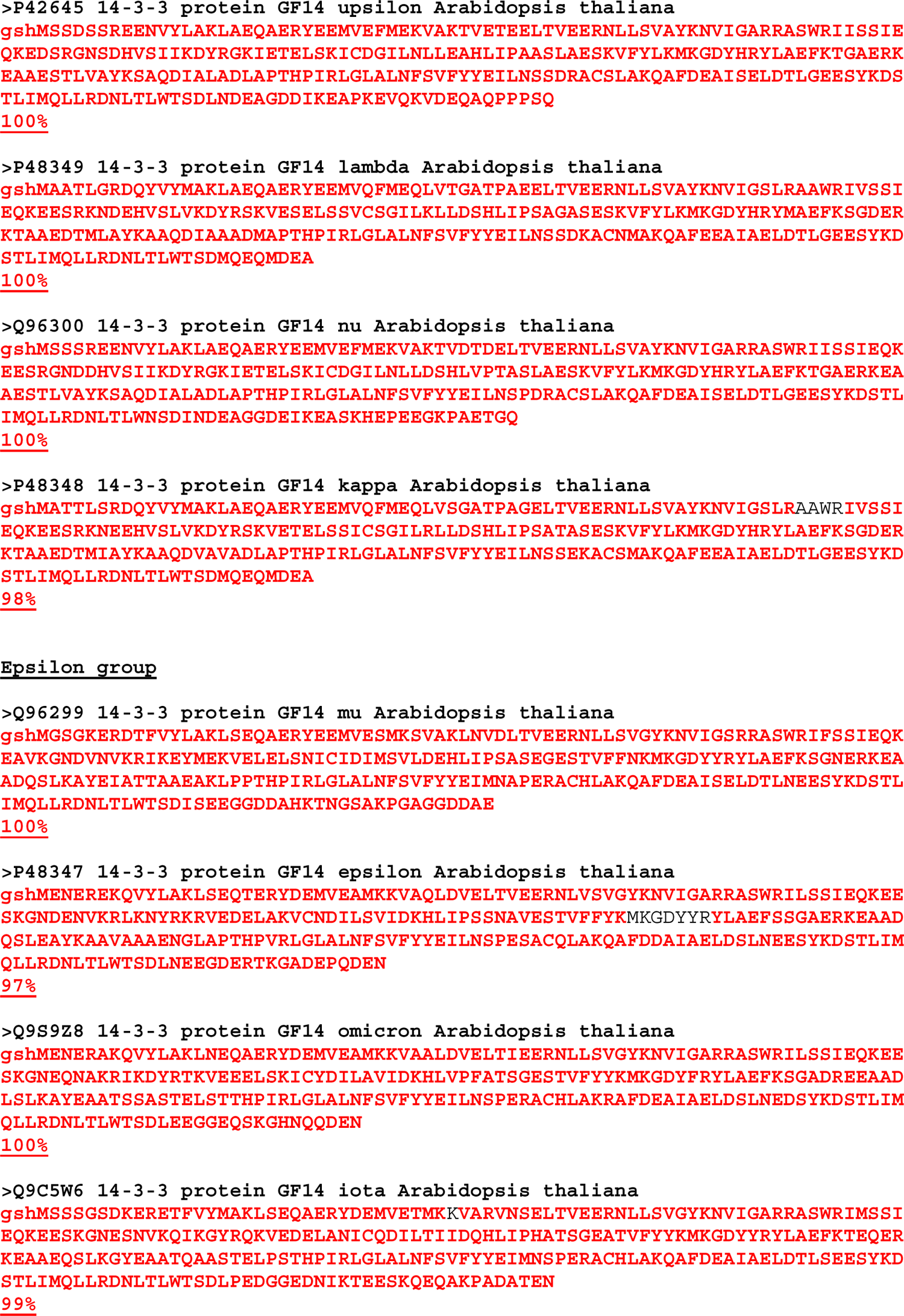
*A. thaliana* 14-3-3 isoforms studied in this work. **A**. Percent identity matrix showing the homology between 13 isoforms of *A. thaliana* and between each of them and human 14-3-3 zeta (ζ). Percentage of identity is color coded from low (red) to high (green). Note that pi isoform is a very distinct plant 14-3-3 isoform. **B**. Alphafold model of 14-3-3 iota homodimer showing the location of the mutations Thr-to-Phe (M1), Asn-to-His (M2) and Gly-to-Ala (M3) mimicking residues conserved in non-epsilon isoforms. **C**. Alignment of 13 *Arabidopsis* 14-3-3 isoforms and human 14-3-3 zeta performed in MCoffee (Wallace *et al*., 2006) using default parameters and visualized using jalview (Procter *et al*., 2021). The boxes are colored by sequence identity using blue shading. Positions M1 (Thr32), M2 (Asn84) and M3 (Gly54) selected for mutation in 14-3-3 iota are indicated by arrows. **D.** Amino acid sequences of 14-3-3 isoforms from *A. thaliana* studied in this work. Small letters represent vector-derived residues, *A. thaliana* 14-3-3 protein sequences are shown with capital letters. Red letters indicate parts of the sequence confirmed by fingerprint MALDI MS. Black letters indicate residues not covered by MALDI MS. Uniprot identifiers are presented for each isoform in the beginning of the title. Bold red underlined numbers after each sequence represent MALDI MS percent of the sequence coverage.

**Fig. S2.**
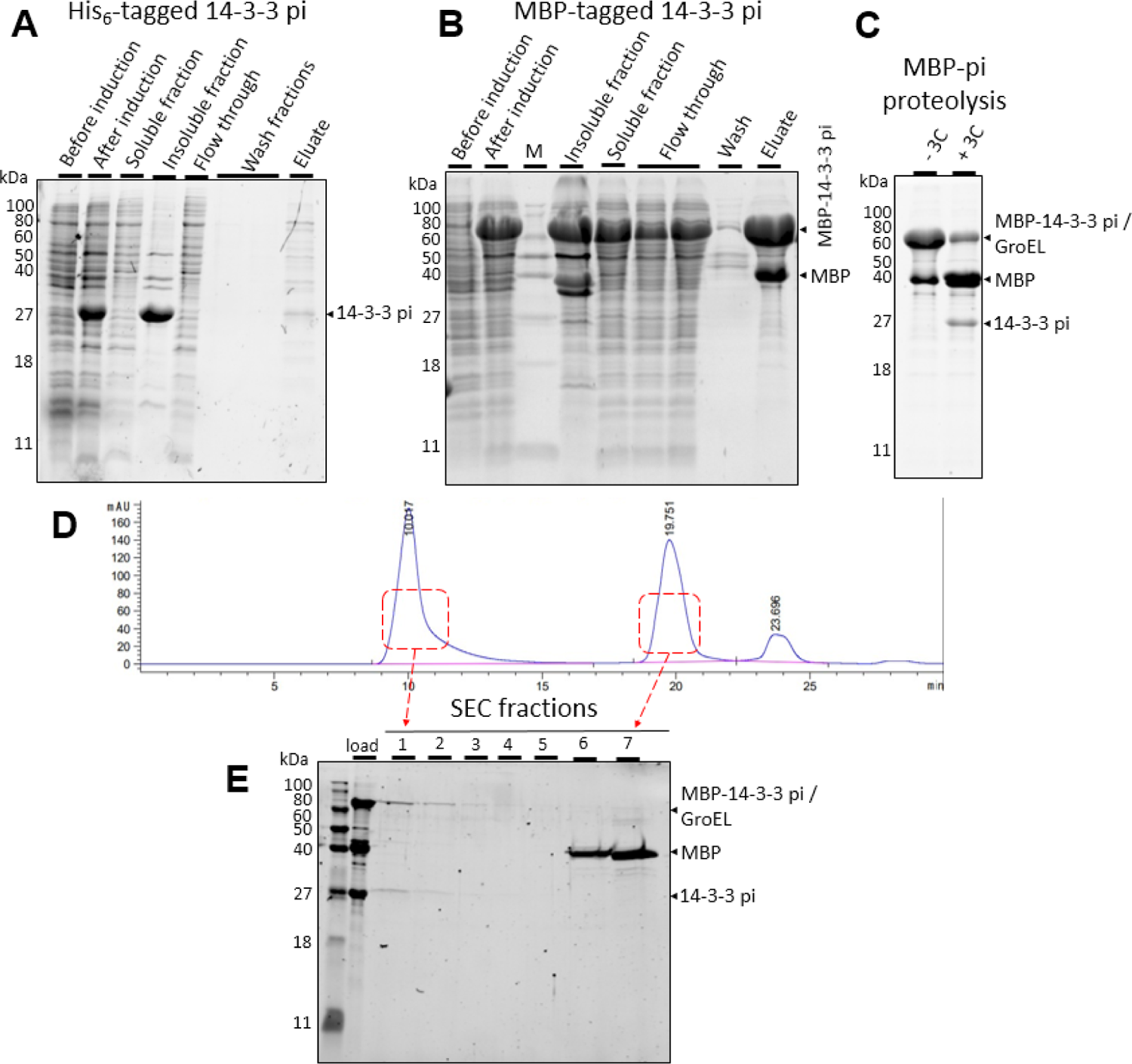
Attempts to obtain recombinant 14-3-3 pi from *A. thaliana*. **A**. Extraction and IMAC of 14-3-3 pi expressed as an N-terminally His-tagged variant. Notwithstanding the large expression yield, the predominant amount of protein is found in the pellet after cell lysis. The eluate from IMAC contains only traces of 14-3-3 pi band which is severely contaminated by other proteins. **B**. Expression and MBPtrap chromatography of 14-3-3 pi fused with the cleavable N-terminal MBP tag. The significant solubilization of 14-3-3 pi by MBP is notable. **C.** MBP-14-3-3 pi cleavage by 3C protease leads to the release of MBP and the target protein. The remaining band at ∼60 kDa can not be cleaved even at 3C excess, and likely corresponds to an endogenous chaperone. **D**. SEC profile of the cleaved MBP 14-3-3 pi preparation showing that the main fraction of the target protein is aggregated and elutes in column void volume. Superdex 200 Increase 10/300 column was used, 0.8 ml/min flow rate. **E**. SDS-PAGE analysis of the SEC fractions demonstrating the coelution of 14-3-3 pi and the 60 kDa band tentatively corresponding to a chaperone protein. MBP elutes at the end of the SEC profile.

**Fig. S3.**
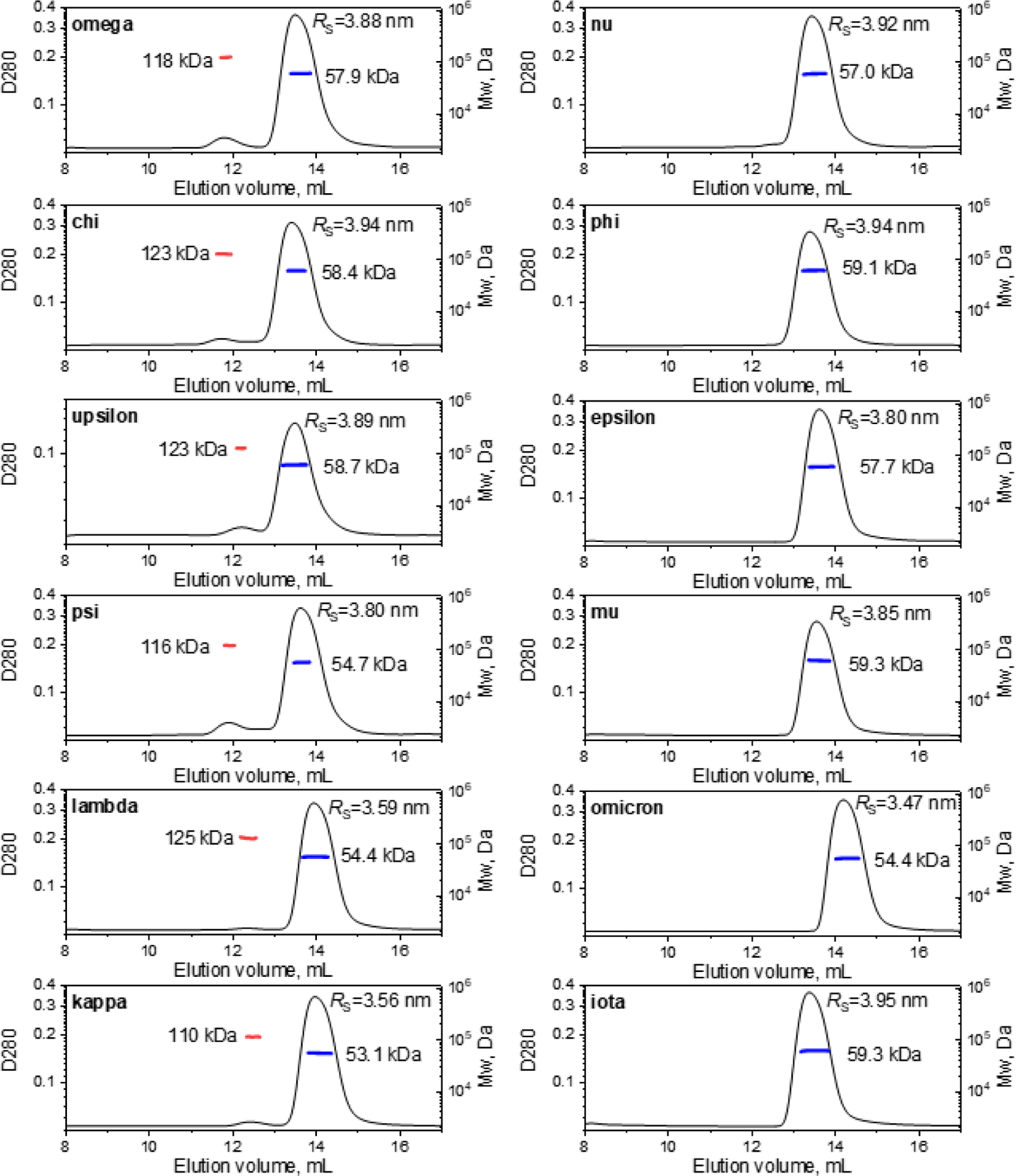
*A. thaliana* 14-3-3 isoforms assemble into dimers at 100 μM protein sample concentration. SEC-MALS profiles for each isoform obtained from a Superdex 200 Increase 10/300 column at 0.8 ml/min are shown along with the absolute *M*_w_ distributions across the main dimeric peak (blue lines) and the corresponding average *M*_w_ value. Note that the six isoforms presented on the left column display also a small peak of tetramers (red *M*_w_ distributions). Stokes radii (*R*_S_) of 14-3-3 peaks were calculated from column calibration. All samples were analyzed prior to preparative SEC purification to reveal all oligomeric forms potentially formed.

**Fig. S4.**
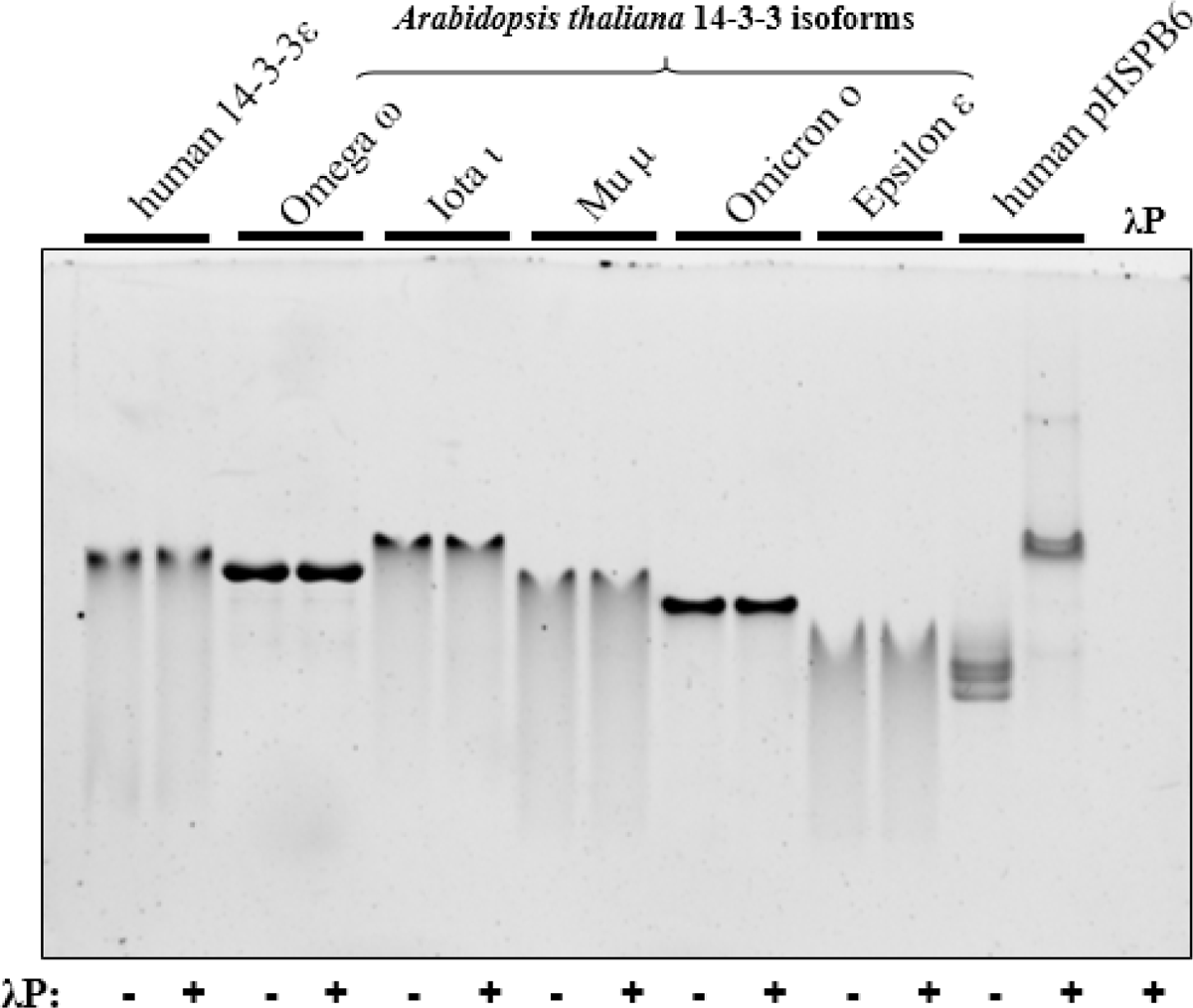
Dephosphorylation of recombinant 14-3-3 samples by lambda phosphatase (λP). The final purified samples of 14-3-3 isoforms from *A. thaliana*, human 14-3-3 epsilon were subjected to dephosphorylation reaction catalyzed by λP (5 h at 20 °C). Positive control was represented by dephosphorylation of human small heat shock protein HSPB6 pre-phosphorylated by PKA (Sluchanko *et al*., 2017; Tugaeva *et al*., 2023). λP at an equivalent amount was loaded on the last lane. Note that under the conditions leading to a complete dephosphorylation of HSPB6, as judged from its discrete upward shift on native-PAGE, no changes of the 14-3-3 bands were observed, strongly disfavoring the hypothesis that the monomerization of epsilon-group *Arabidopsis* 14-3-3 isoforms is associated with their inadvertent phosphorylation.

**Fig. S5.**
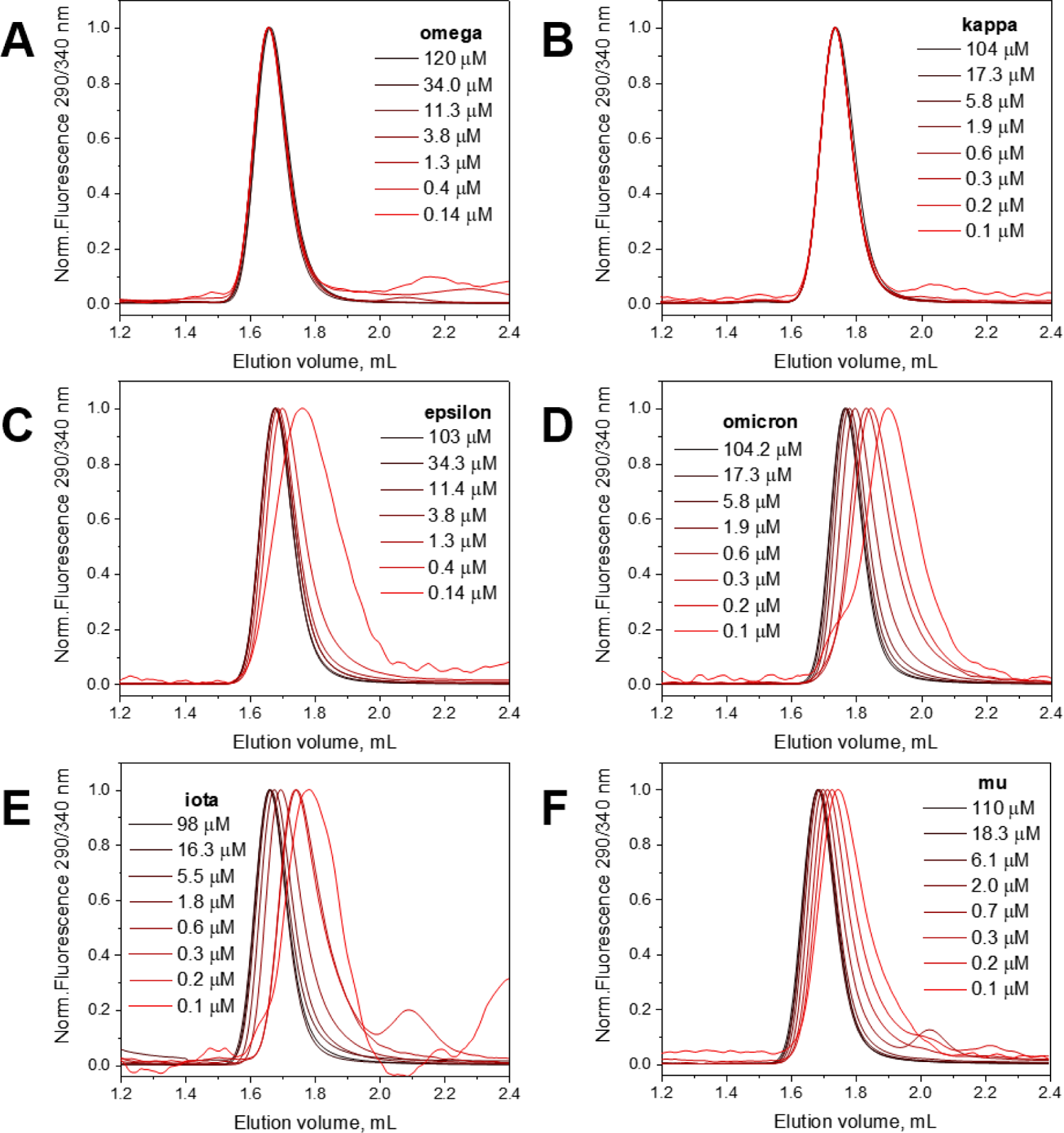
All epsilon group isoforms show concentration-dependent 14-3-3 dimer dissociation. **A-F**. Analytical SEC profiles of 14-3-3 omega (A), kappa (B), epsilon (C), omicron (D), iota (E) and mu (F) at different protein concentrations in the loaded sample (indicated in μM). Elution profiles from a Superdex 200 Increase 5/150 column (0.45 ml/min) were followed by Trp fluorescence detection (excitation 290 nm, emission 340 nm).

**Fig. S6.**
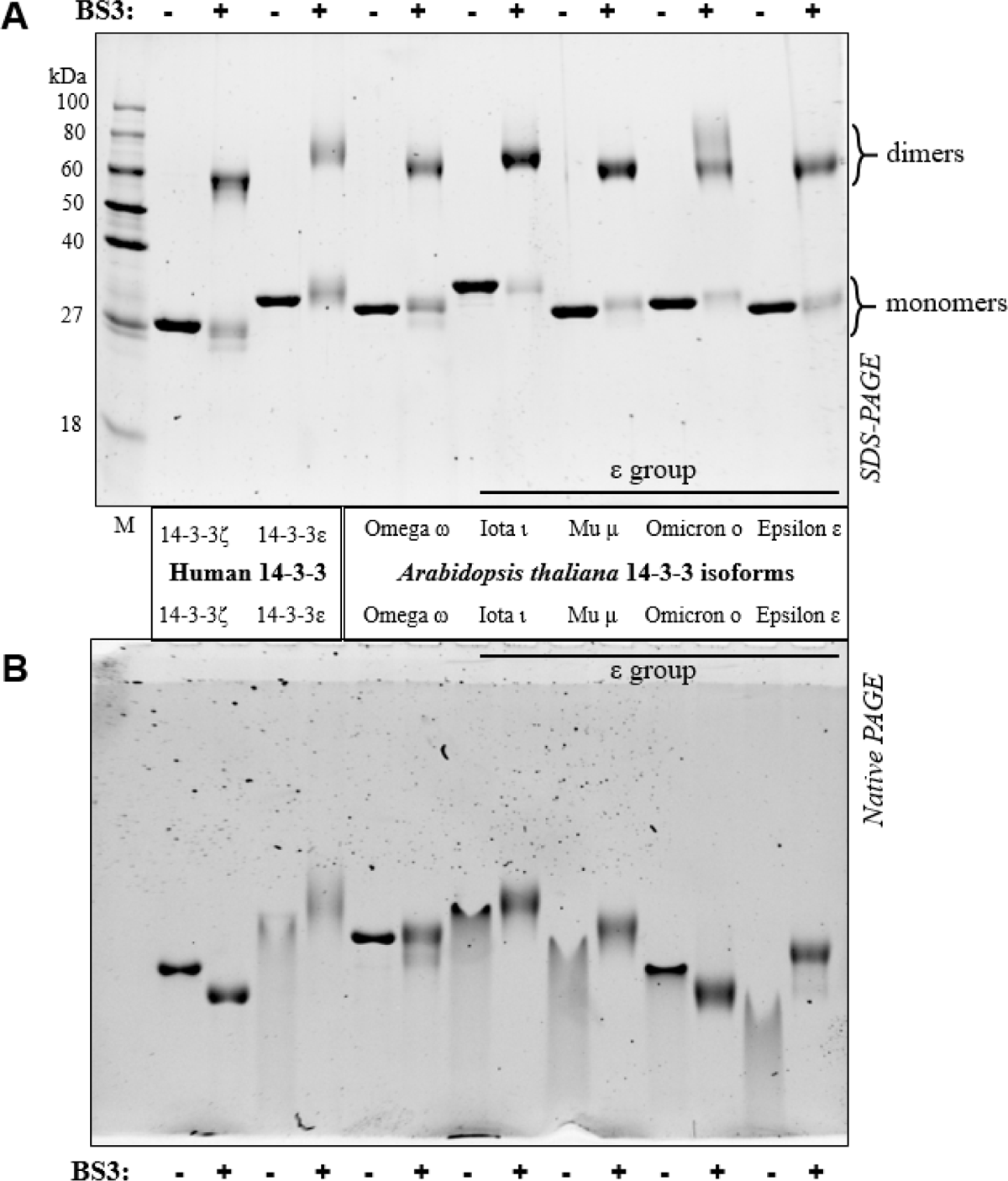
Both non-epsilon and epsilon group *Arabidopsis* 14-3-3 isoforms form dimers upon chemical crosslinking with BS3. Indicated plant and human 14-3-3 isoforms were incubated in the absence (-) or in the presence (+) of 0.7 mM BS3 crosslinker (final concentration) for 40 min at 20 °C and subjected to SDS-PAGE (12.5% acrylamide) (**A**) and native PAGE (15% acrylamide, 2 h runtime) (**B**). 0.4 μg of each protein was loaded on the gels. Note that BS3 crosslinking fixed 14-3-3 proteins in the dimeric state, without causing the formation of any non-specific higher-order oligomers. On native gel, the crosslinked epsilon-group *Arabidopsis* 14-3-3 isoforms (iota, mu and epsilon) focused and lifted, apparently approaching the position of the fully assembled dimers. The same scenario was observed for human 14-3-3 epsilon, whereas *Arabidopsis* 14-3-3 omicron and human 14-3-3 zeta behaved differently in that their bands moved down upon crosslinking, and the smearing of omicron disappeared. This may indicate that BS3-related chemical modification of some lysines in 14-3-3 can make the molecule more compact, for instance, due to preventing some expanded conformations.

**Fig. S7.**
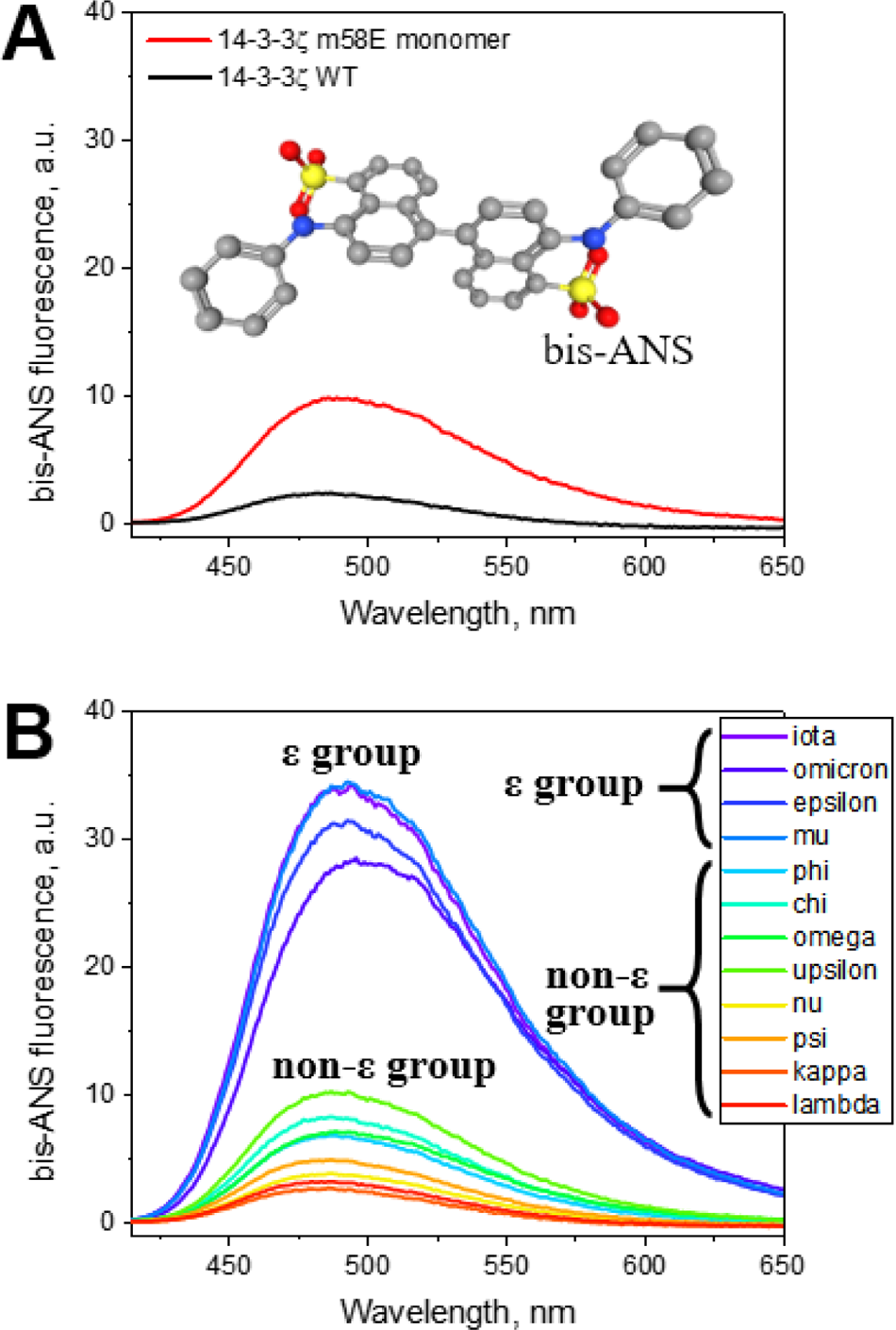
Surface hydrophobicity of 14-3-3 proteins probed by bis-ANS. **A**. Steady-state bis-ANS fluorescence spectra recorded at 25 °C after incubation of the dye (10 μM) with either human 14-3-3ζ wild-type or the monomeric mutant thereof (2 μM each). Bis-ANS structure is shown as an insert. **B**. Steady-state bis-ANS fluorescence spectra recorded after incubation of the dye (10 μM) with each of the twelve 14-3-3 isoforms from *A. thaliana* (2 μM each) under the same parameters as were used for obtaining the spectra shown on panel A. Note the variability within the epsilon and non-epsilon groups: for example, the difference between the monomer and dimer of human 14-3-3 zeta was in the same range as the difference between non-epsilon isoforms upsilon and psi.

**Fig. S8.**
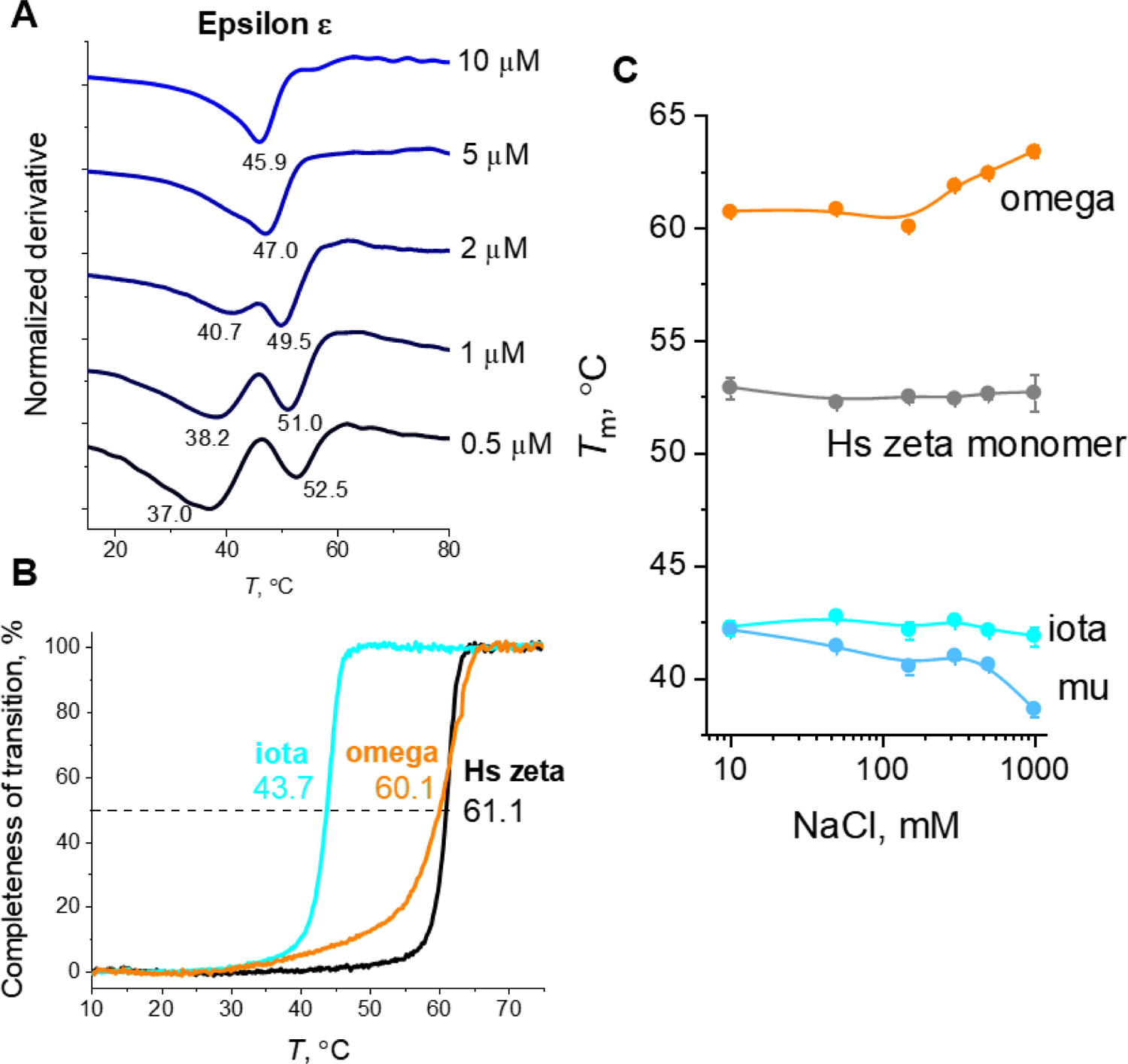
**A.** Thermal stability of *Arabidopsis* 14-3-3 epsilon studied by DSF at 5 different protein concentrations. The amount of ProteOrange dye was the same (5X). Buffer −20 mM HEPES pH 7.5, 150 mM NaCl. The samples were pre-equilibrated at 10 °C and then heated to 90 °C at a heating rate of 1 °C/min. **B**. Validation of the thermofluor data using intrinsic Trp fluorescence intensity for iota and omega. Human 14-3-3 zeta wild-type (Hs zeta) was used as control. Final protein concentration in all cases was 5 μM. Heating from 10 to 80 °C (1 °C/min). **C**. Effect of ionic strength on *T*_m_ values determined by thermofluor for the selected 14-3-3 isoforms (2 μM).

**Fig. S9.**
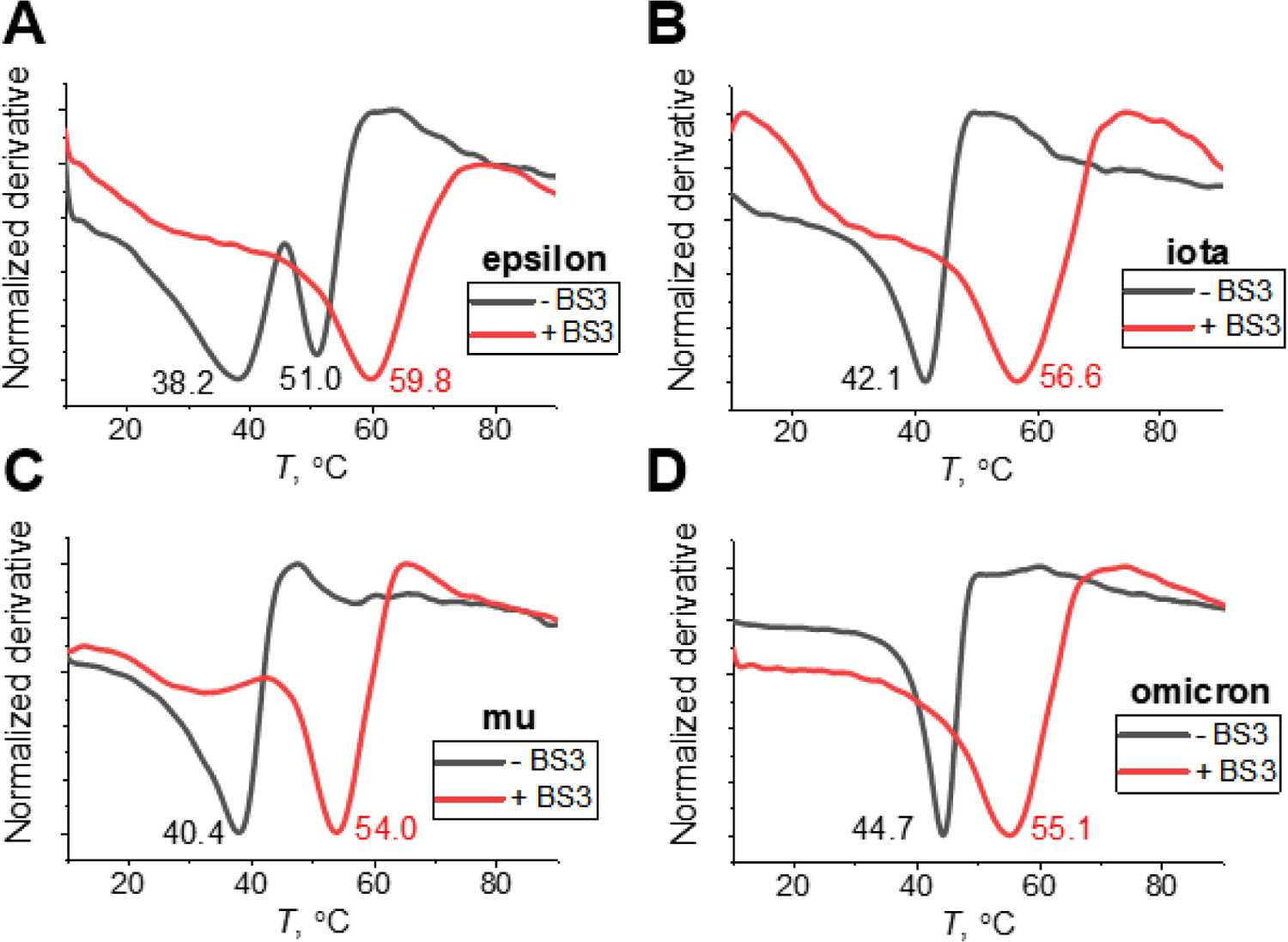
Thermofluor data for uncrosslinked and BS3-crosslinked *Arabidopsis* epsilon-group 14-3-3 isoforms epsilon (A), iota (B), mu (C) and omicron (D). Final protein concentration in all cases was 1 μM. Buffer −20 mM HEPES pH 7.5, 150 mM NaCl. The samples were pre-equilibrated at 10 °C and then heated to 90 °C at a heating rate of 1 °C/min.

**Fig. S10.**
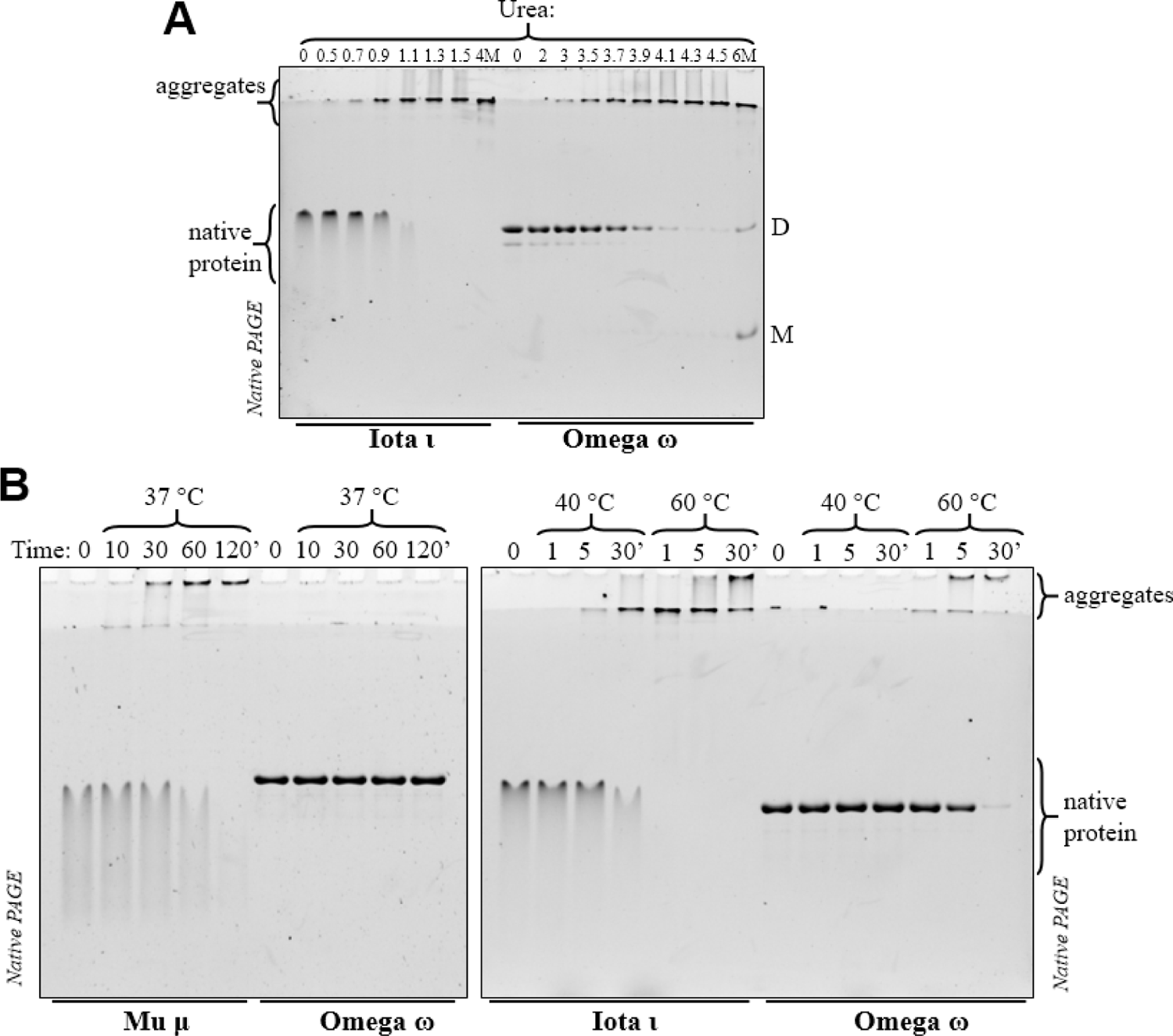
Epsilon-group *Arabidopsis* 14-3-3 isoforms show aggregation propensity and susceptibility to proteolytic degradation. **A**. Urea-induced denaturation of iota and omega analyzed by following disappearance of native protein at increasing urea concentration. Protein concentration was 0.5 mg/ml. D and M indicate positions of omega dimers and monomers after refolding during the run. **B**. Kinetics of aggregation at 37, 40 and 60 °C reveals markedly different stability of epsilon and non-epsilon group 14-3-3 isoforms of *A. thaliana*. Protein samples (35 μM) were incubated at indicated temperature for a given period of time and then loaded on native-PAGE. As a result of protein denaturation and aggregation proteins are partitioned between native and aggregated forms shown by brackets on the right.

**Fig. S11.**
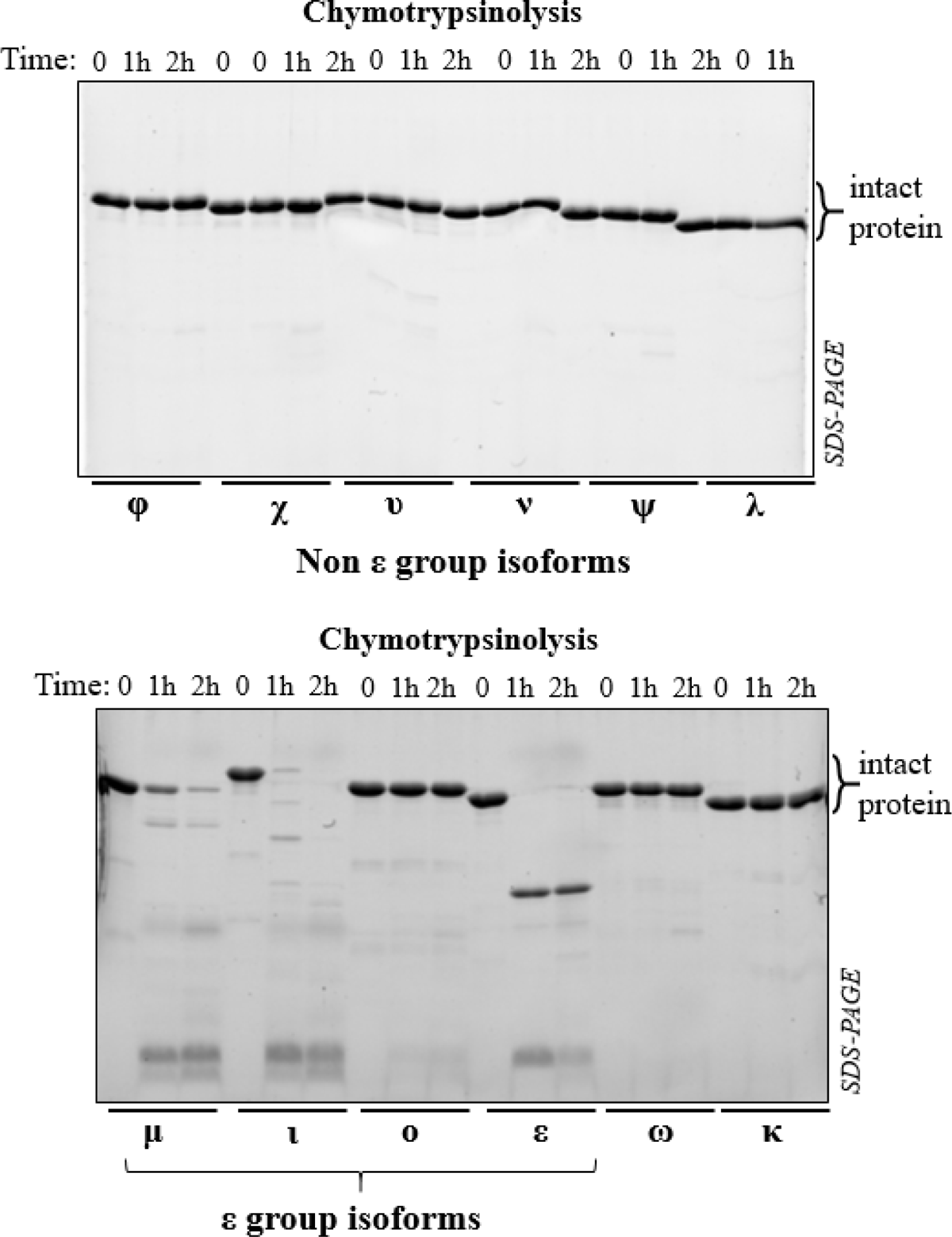
Limited chymotrypsinolysis of *Arabidopsis* 14-3-3 isoforms at a substrate:protease weight ratio of 200:1 at 27 °C. Protein concentration was 0.5 mg/ml. Note that none of non-epsilon group isoforms degraded significantly under these conditions.

